# Asymmetric patterns of nucleosome positioning sequences in protein-coding regions

**DOI:** 10.1101/2023.04.16.537090

**Authors:** Hiroaki Kato, Mitsuhiro Shimizu, Takeshi Urano

**Author notes:** To whom correspondence should be addressed. Hiroaki Kato Department of Life Sciences, Shimane University School of Medicine 89-1 Enya-cho, Izumo, Shimane 693-8501, Japan.

## Abstract

Although nucleosome positioning is key to epigenetic regulation, how the DNA sequence contributes to positioning remains elusive, especially in the context of transcription direction. Analysis of nucleotide bases with respect to the nucleosomal DNA coordinates requires precise nucleosomal mapping information on the genome. However, currently available base-pair-resolution nucleosome maps based on cysteine-mediated chemical cleavage do not fully satisfy the requirement due to method-specific cleavage biases. Here, we generated a chimeric nucleosomal DNA model to achieve less-biased prediction. The model revealed that yeast protein-coding sequences have higher affinity for the promoter-proximal half of nucleosomes than for the distal half. Strikingly, peaks of calculated affinity scores for the promoter-proximal half periodically matched the first few nucleosome positions. Detailed analysis of nucleotide bases revealed that the AA dinucleotide in the left side of the top strand contributes to nucleosome detection frequency in intergenic regions, while the complementary dinucleotide TT is preferred in the other side. In contrast, the sense strand is AA-rich throughout the nucleosome coordinate in protein-coding regions, which is consistent with asymmetric affinity. These data suggest that eukaryotes have evolved DNA sequences with asymmetric affinity for nucleosome formation to maintain epigenetic integrity of protein-coding regions.

## INTRODUCTION

The nucleosome, in which ∼147 bp DNA wraps around a histone octamer (1,2), is the fundamental unit of chromatin and is vital for epigenetic regulation (3,4). Besides those in intergenic regions, nucleosomes are formed with regular spacing in protein-coding genes (5,6). The regularity of nucleosome arrays relies on chromatin remodeling factors and histone chaperones accompanying RNA polymerase II (6–10). In addition to the functions of *trans*-acting factors, the DNA sequence *per se* is thought to have an intrinsic ability to help determine nucleosome positioning despite the lack of detailed studies in terms of transcription direction (11–14). Recent structural and molecular dynamics simulation studies have uncovered the dynamic nature of nucleosomes and provided insights in how genetic information stored in nucleosomes is decoded efficiently (15–20). However, as these studies utilize particular sequences that are favorable for *in vitro* nucleosome positioning, understanding of the impact of natural DNA sequences on *in vivo* positioning, especially in the context of transcription, is deficient.

Determination of nucleosome positions along the genome is key to studying DNA sequences with respect to nucleosomal coordinates, which consist of 147 nucleotide positions (-73 to +73) with a dyad base located at the midpoint (position = 0). Early genome-wide studies utilized micrococcal nuclease (MNase) to map *in vivo* nucleosomes; however, low mapping resolution was obtained due to the sequence preference of the enzyme (12,21). Higher resolution nucleosome mapping was achieved by means of cysteine-mediated chemical cleavage of DNA with histone H4-S47C or H3-Q85C mutations (11,21–23). As the hydroxyl radical produced in the localized Fenton reaction attacks the deoxyribose ring (24,25), it was formerly expected that the chemical cleavage was not affected by the local sequence of nucleotide bases around the cleaved sites. Contrary to expectations, prominent occurrence of some nucleotide bases near H4-S47C’s target sites (11) and difference in dinucleotide patterns between the nucleosomes called by the H4-S47C and H3-Q85C methods (21) have implied that the chemical mapping methods have their own sequence biases in nucleosome calling. However, the potential biases have not yet been clarified, due to the lack of appropriate methods for classifying short DNA sequences in terms of appropriateness for nucleosome formation or nucleosome calling.

We recently reported that chemical map–based statistical models performed better than classic MNase-seq–based models in predicting nucleosomal translational positions and rotational settings (26). Specifically, we demonstrated that the appropriateness of given 147-bp sequences for nucleosome formation can be evaluated with histone binding affinity (HBA), as theoretically proposed by Xi et al. (27), when the calculation was based on chemical maps. Chemical-map–based HBA scoring has been utilized for assessment of natural and artificial sequences (9,26,28). We also developed a method to evaluate the appropriateness of 20 to 21-bp subsegments of nucleosomal DNA for being located at particular positions in the nucleosomal coordinate (**Figure 1A**, Local HBA segments), and the calculated affinity scores for tested sequences agreed well with the literature (26,29). However, as the chemical map–based models used in these studies were based on the *in vivo* nucleosome maps determined with H4-S47C mutations, a concern could be raised that the prediction *per se* is also affected by the as yet poorly studied biases in DNA cleavage and subsequent nucleosome calling. Even if true, the models can be utilized to characterize the sequence dependent biases in chemical mapping.

**Figure 1.**
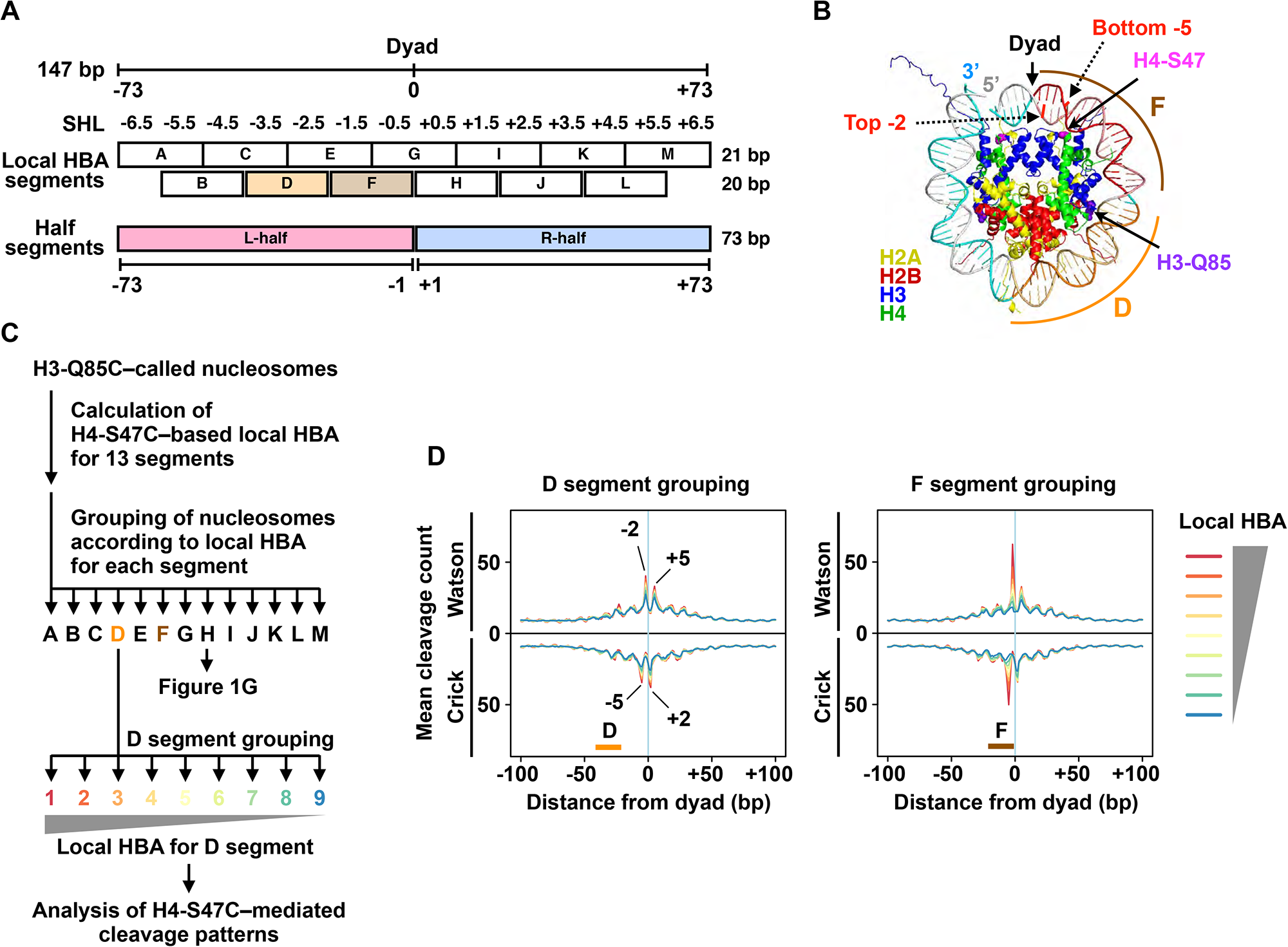
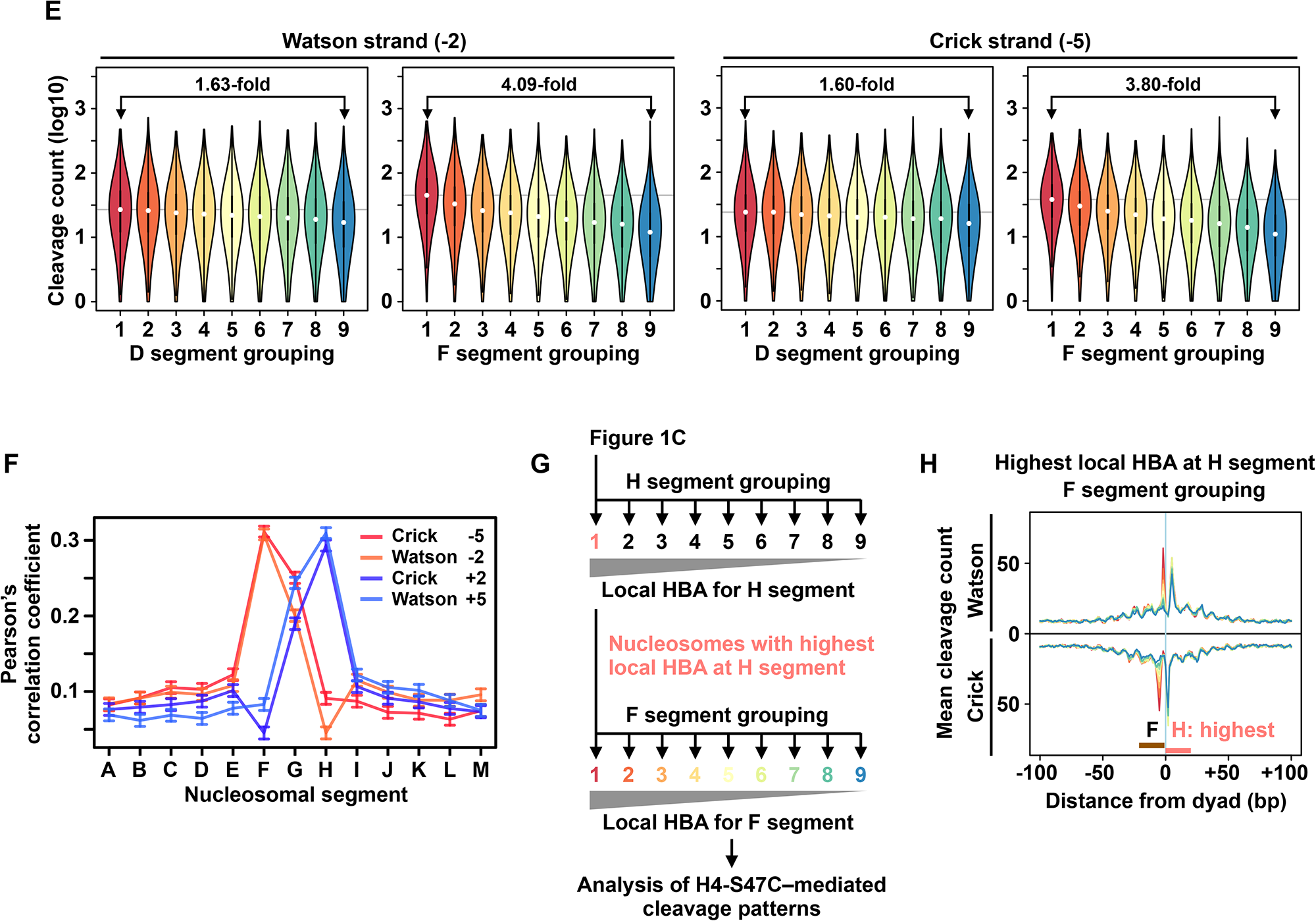
Sequence-dependent biases in H4-S47C-mediated DNA cleavage. (**A**) Nucleosomal DNA segments analyzed in this study. (**B**) Sites of H4-S47C-mediated cleavage. Double stranded DNA wrapped on the front side of the nucleosome (1AOI) is shown. The -2 and -5 nucleotide positions in the top (light gray or lighter) and the bottom (light blue or darker) strands, respectively, are shown. D and F segments are colored in the shades of orange and brown, respectively. (**C**) Scheme of nucleosome grouping according to local HBA scores. Grouping was performed for all thirteen segments (A to M). (**D**) Mean cleavage count along the nucleosomal coordinate for each local HBA group. Lines for lower local HBA groups are overlaid on those for higher score groups. Locations of the segments used for local HBA calculation and nucleosome grouping are indicated as horizontal bars at the bottom. Nucleotide positions of cleavage peaks are indicated. (**E**) Cleavage counts at particular nucleotide positions in each local HBA group. Median cleavage counts for the groups with highest local HBA scores are indicated as horizontal lines. Lines of cleavage counts for lower local HBA groups are overlaid on those of higher local HBA groups. (**F**) Pearson’s correlation coefficient between cleavage counts at indicated positions and local HBA at indicated segments. 95% confident intervals are shown. (**G**) Scheme of secondary nucleosome grouping. For the primarily selected nucleosomes with highest local HBA at segment F or H, secondary grouping was performed for all thirteen segments. (**H**) Mean cleavage count along the nucleosomal coordinate as in **D** for selected nucleosomes with highest local HBA at H segment. See **D** for the color key. (**D** and **H**) For the results of the other groupings, see **Supplementary Figures S1, S2,** and **S3**.

In the present study, we applied chemical map–based models to evaluate the effect of local DNA sequence on cleavage efficiency, thereby verifying that the chemical mapping methods indeed have sequence preferences in nucleosome calling. With an ideal nucleosome model generated from a set of chimeric nucleosomal DNA sequences that exclude the biased elements, we discovered an unexpected property of DNA sequences in yeast protein-coding regions. With higher AA dinucleotide composition in the sense strand, DNA sequences that are appropriate for wrapping in the promoter-proximal half of the nucleosome are embedded in coding sequences in a periodic manner that facilitates the determination of nucleosome positions. Collectively, our findings provide insight into how DNA sequences contribute to nucleosome dynamics in the context of transcription, especially in protein-coding regions.

## MATERIAL AND METHODS

### Computational environment

The GNU R statistical environment (https://www.r-project.org/, ver. 4.1.1) and its built-in functions were used for most of the study, unless otherwise stated. DNA sequences were obtained from the yeast genome sacCer3 (R64) and the mouse genome GRCm38 (mm10) with the functionalities of Biostrings (https://doi.org/doi:10.18129/B9.bioc.Biostrings, ver. 2.60.2). Heatmaps were drawn with the levelplot function of rasterVis (https://cran.r-project.org/package=rasterVis, ver. 0.51.2). Nucleosome structures were drawn using MacPyMOL (v1.8.4.1).

### Preparation of published nucleosome datasets

Dyad positions of yeast representative nucleosomes called by the H4-S47C method (11) <u>lifted over</u> the sacCer3 coordinate (30) were used as an initial set of H4-S47C–called nucleosomes (n = 67,548). Three bigWig format files from the H3-Q85C study (21) with accession identifiers GSM2561057, GSM2561058, and GSM2561059 from the GEO database (https://www.ncbi.nlm.nih.gov/geo) were converted to the Wig format with UCSC bigWigToWig (ver. 377). The number of 51 ± 7 bp nucleosomal fragments assigned to individual dyad positions (hereafter designated as “dyad scores”) were combined to obtain a normalized genome-wide map of H3-Q85C–called nucleosomes, as described (21). We defined the NCP scores for H4-S47C–called nucleosomes (11) and the above-mentioned dyad scores for H3-Q85C–called nucleosomes (21) as the method-specific nucleosome detection frequency.

### Selection of representative nucleosomes for sequence analyses

Nucleosomes were processed according to the detection frequency of individual dyads assigned to given genomic coordinates. By identifying nucleosome dyads with the highest score in the surrounding ± 53 bp window, the H3-Q85C–called whole genomic representative nucleosomes were selected (n = 60,612). For the H4-S47C–called nucleosome dyads, dyads not located within 300 bp of the chromosome ends were selected as being representative (n = 67,510). The 147-bp Watson strand sequences in which the selected dyads were located at the midpoint were obtained as the top strands of nucleosomal DNA for the representative nucleosomes. From the H3-Q85C–called and H4-S47C–called whole genomic representative nucleosomes, those not overlapping with protein-coding genes were selected as intergenic nucleosomes (n = 11,158 and 12,130, respectively).

For nucleosomes found in protein-coding regions (hereafter denoted as coding nucleosomes), the first and following representative nucleosomes that overlap with coding sequences were defined as +1, +2, and +3 coding nucleosomes, and so on. Coding nucleosomes with at least one additional representative nucleosome in the same gene and transcribed unidirectionally were used for the analyses. The sense strand of the gene was treated as the top strand of the nucleosome. Results for the +2 nucleosomes were selected for the main figures as they represented sequence patterns generally found in the tested nucleosomes.

### Symmetrization of nucleosome datasets

Dinucleotide analyses of the whole genomic and intergenic nucleosomes were conducted by combining forward sequences with the reverse complement sequences to obtain averaged, less biased results. The detection frequency for each nucleosome was assigned to both the forward and reverse complement sequences. Each of the forward and reverse complement nucleosomes were regarded as distinct single nucleosomes in the symmetrized dinucleotide analyses. Note that symmetrization was not applied for coding nucleosomes as it would prevent analysis regarding transcription direction.

### Construction of native and chimeric nucleosome models

Of the 67,548 H4-S47C–called nucleosome dyads, 22 were located within 200 bp from the chromosomal ends, and 174 did not have any H3-Q85C–called nucleosome dyads in the surrounding ± 50 bp regions. In order to construct statistical models, we selected a set of representative H3-Q85C–called nucleosomes as those with the highest dyad scores in the ± 50 bp window of each H4-S47C–called nucleosome dyad. As a result, an equal number of representative nucleosomes (n = 67,352) was selected for each chemical mapping method. These matched representative nucleosomes were used for statistical model construction. A set of chimeric nucleosomal DNA sequences was created by concatenating the central 33-bp sequences (-16 to +16) from the H3-Q85C–called nucleosomes and the outer sequences (-73 to -17 and +17 to +73) from the H4-S47C–called nucleosomes. Each chimera was created from a pair of H4-S47C–called and H3-Q85C–called nucleosomes that were located at the nearest chromosomal positions. Fourth-order time-dependent matched nucleosome models were constructed from the native (i.e., H3-Q85C–called and H4-S47C–called) and chimeric nucleosomal sequences (n = 67,352 for each) as previously described (26,27). For presenting dinucleotide patterns of nucleosomal sequences, frequencies of selected dinucleotides (AA, AT, TA, TT, CC, CG, GC, and GG) starting at each nucleotide position in the nucleosomal coordinate were calculated. Frequencies of WW and SS dinucleotides, in which W and S refer to a “weak” base (A or T) and a “strong” base (C or G), were calculated as cumulative frequencies of corresponding dinucleotides.

### Calculation of histone binding affinity

HBA was defined by Xi et al. (27) as the log likelihood ratio of a given 147-bp sequence to be in a nucleosome state versus a linker state. Thus, the HBA score for a 147-letter sequence *x* centering at nucleotide position *i* (*a_i_*) is,

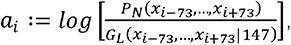

in which *P_N_* is a time-dependent fourth-order nucleosome model and *G_L_*is a homogeneous fourth-order linker model (27). We split the 147-bp nucleosomal DNA into left (nucleotide positions -73 to -1) and right (+1 to +73) segments to calculate HBA scores for each segment (**Figure 1A**, Half segments). Like the full length HBA calculation, the HBA score for the left (L-half) segment of a 147-bp region *x* centering at position *i* (*L_i_*) is,

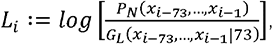

and that for the right (R-half) segment (*R_i_*) is,

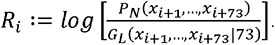

Therefore, the sum of HBA scores for the L- and R-halves nearly equals the whole nucleosomal HBA for a given 147 bp sequence. Both the top and bottom strand sequences are considered in the HBA calculation (27). Thus, it can be said that the HBA score for a 147-letter sequence will be the same as that for the reverse complement sequence, and that L- and R-half segments for a palindromic 147-bp sequence will have the same HBA scores. Each of the three nucleosome models described above was used to calculate the probability of a given sequence being a nucleosome or part thereof. For the linker model, the model previously constructed from the map of H4-S47C–called nucleosomes was used (26). Local HBA scores (26) were calculated in the same manner. HBA analysis of the Widom 601 sequence was performed as described (26) with the H4-S47C–based and chimeric models constructed in the present study. The Fortran code for HBA calculation from NuPoP (ver. 1.20.0) (27) was adapted for these purposes as previously done (26). Note: A newer version of NuPoP (ver. 2.4.0, https://doi.org/doi:10.18129/B9.bioc.NuPoP) is available for duration hidden Markov model–based predictions.

### Grouping of nucleosomes according to local HBA scores or detection frequency

For each representative nucleosome, local HBA scores for 13 segments, namely A to M, were calculated with the statistical models constructed in the present study unless otherwise stated. The nucleosomes were evenly grouped into nine groups according to the order of local HBA scores for each segment or nucleosome detection frequency. From the set of tested nucleosomes, those with the top 1% and bottom 10% detection frequencies were excluded as exceptional cases. This process omitted the nucleosomes in repeated sequences and those called despite low cleavage evidence.

### Analyses of H4-S47C–mediated cleavage counts on the nucleosomal coordinate

H4-S47C–mediated cleavage counts from Henikoff et al. (SRX372005, SRX372006, SRX372007) (31) were obtained as previously described (30). The nucleotide positions 1 bp upstream of the 5’-end of the mapped reads were regarded as cleavage sites. For each of the H3-Q85C–called representative nucleosomes (n = 60,612), H4-S47C–mediated cleavage counts in the surrounding ± 101 region of the Watson and Crick strands were stored. Here, the Watson and Crick strands of the genome correspond to the top and bottom strands of the nucleosome structure, respectively. Grouping of nucleosomes according to local HBA scores was performed as described above using the *localHBA* function (species = “sc”) of nuCpos (26). Mean cleavage counts along the nucleosomal coordinate were plotted for the grouped nucleosomes. For each nucleosome, cleavage counts at the -2 and +5 positions of the Watson strand and those at the -5 and +2 positions of the Crick strand were obtained. Pearson’s correlation tests were performed between these cleavage counts and local HBA scores for each segment. For the analysis of nucleosomes with the highest local HBA scores for the F or H segments (n = 6,735 each), secondary grouping of nucleosomes was performed according to the local HBA scores for other segments.

### HBA analysis of protein-coding sequences

For HBA analysis of yeast protein-coding genes, those longer than 1.5 kb and harboring the +1 nucleosomal dyad determined by Chereji et al. (21) were selected (n = 2,148). Sense strand sequences from the nucleotide positions -573 to +1073 relative to the +1 dyad were utilized for calculation of HBA scores. In order to analyze murine protein-coding sequences, the NCP scores of Voong et al. (23) were remapped from the mm9 to mm10 coordinates. For the mm10 coordinate, a full set of “RefSeq Select” transcripts was available (https://www.ncbi.nlm.nih.gov/refseq/refseq_select/). The transcript dataset contained 2,656 >500 bp exons with no untranslated regions, in which 1,606 had a representative nucleosome determined by the H4-S47C method (23) within the central 147 bp region. The exons had at least 177 bp coding sequences from the central nucleosome dyad. Sense strand sequences from the nucleotide positions -573 to +573 relative to the central dyad were utilized for HBA calculation. For both the yeast and mouse sequences, full length, R-half, and L-half HBA scores for each possible 147 bp sequence were assigned to the respective dyad position. Mean HBA scores were smoothed using a 51-bp centered moving average.

### Evaluation of the impact of dinucleotides at distinct positions on nucleosome detection frequency

Pairs of dinucleotides to be tested were defined as follows: AA versus TT (AA/TT), AG/CT, GA/TC, GG/CC, and RR/YY, in which R stands for one of the two purine bases (A and G), and Y stands for p<u>y</u>rimidine bases (C and T). From the whole genomic or intergenic nucleosomes, those containing individual dinucleotides in a pair at a particular position on the nucleosomal coordinate and those containing the corresponding dinucleotide in the pair at the same position were selected. The selection was performed for all possible starting dinucleotide positions (-73 to +72). The Welch t-test was performed to assess the log-transformed detection frequency of the selected nucleosomes for each dinucleotide starting position.

### Dinucleotide frequency rank visualization on the nucleosomal coordinate

For each set of nucleosomes, frequencies of the 16 dinucleotides starting at each nucleotide position were calculated. Each dinucleotide was ranked according to the order of the frequency at its particular position, and presented as a colored bar in two-dimensional spaces. AA and TT dinucleotides were identified by shading the respective bars.

## RESULTS

### Cleavage biases of chemical mapping methods

Previous studies reported that chemical mapping methods with different histone mutations call nucleosomes with different DNA sequence patterns (11,21). Although this suggests that method-specific cleavage biases hinder fair calling of nucleosomes, the influence of the DNA sequence in chemical mapping has not been investigated to date. The suitability of 20 to 21-bp nucleosomal subsequences (called A to M) for nucleosome calling with H4-S47C can be evaluated using local HBA scoring (**Figure 1A**) (26). With this scoring method, we first studied the impact of local DNA sequences on H4-S47C–mediated chemical cleavage. In order to assess whether Cys-mediated chemical cleavage exhibits sequence preference, we analyzed H3-Q85C–called nucleosomes (n = 60,612), which are free from the potential biases of H4-S47C-mediated cleavage (**Figure 1B**). The workflow of the analysis is presented in **Figure 1C**. H4-S47C–based local HBA scores were calculated for each of the H3-Q85C–called nucleosomes. According to the local HBA score for each segment, nucleosomes were evenly grouped into nine groups per segment basis. For each of the 13 segment groupings, H4-S47C–mediated chemical cleavage patterns of Henikoff et al. (31) were then analyzed in relation to the local HBA scores for the segment used for the grouping (**Figure 1D** and **Supplementary Figure S1**).

The left panel of **Figure 1D** shows that cleavage counts around the dyad correlate with HBA scores for the D segment. Indeed, local HBA scores at non-central segments including D exhibited a minor but significant contribution to cleavage around the dyad (Pearson’s correlation test, p < 1x10^-27^, **Supplementary Table S1**). This suggests that the affinity of each DNA segment to histones helps stabilize nucleosomes *in vivo*. In contrast, as exemplified in the right panel of **Figure 1D**, local HBA scores at central segments greatly correlated with H4-S47C–mediated cleavage. For the central F segment grouping, median cleavage counts for the highest local HBA group were 4.09-fold (Watson strand, position -2) and 3.80-fold (Crick strand, position -5) higher than those for the lowest local HBA group (**Figure 1E**, **Supplementary Figure S2** and **Supplementary Table S2**). These values were much higher than those for the D segment grouping: 1.63-fold and 1.60-fold, respectively.

Further, we observed that the sequence around the cleavage sites in one half exerted only a minor contribution to the cleavage in the other half (**Figure 1F**). To clarify this tendency, we performed a secondary grouping of nucleosomes according to local HBA scores from those with the highest local HBA scores for F or H segments (**Figure 1G**). Interestingly, even for the nucleosomes with the highest local HBA scores for the right segment H, cleavage counts in the left side were highly correlated with the local HBA scores for the left segment F (**Figure 1H** and **Supplementary Figure S3**). These observations suggest that the frequency of chemical cleavage can be strongly affected by local DNA sequence around cleavage sites.

Next, we studied the relationship between subsegment sequences and detection frequency of chemically called nucleosomes. Whole genomic H3-Q85C–called and H4-S47C–called nucleosomes were grouped according to their detection frequency (evaluated as dyad scores and NCP scores, respectively), and local HBA scores were analyzed with H3-Q85C–based and H4-S47C–based statistical models (**Figure 2**, left and center). The detection frequency of H4-S47C–called nucleosomes was strongly correlated with

**Figure 2.**
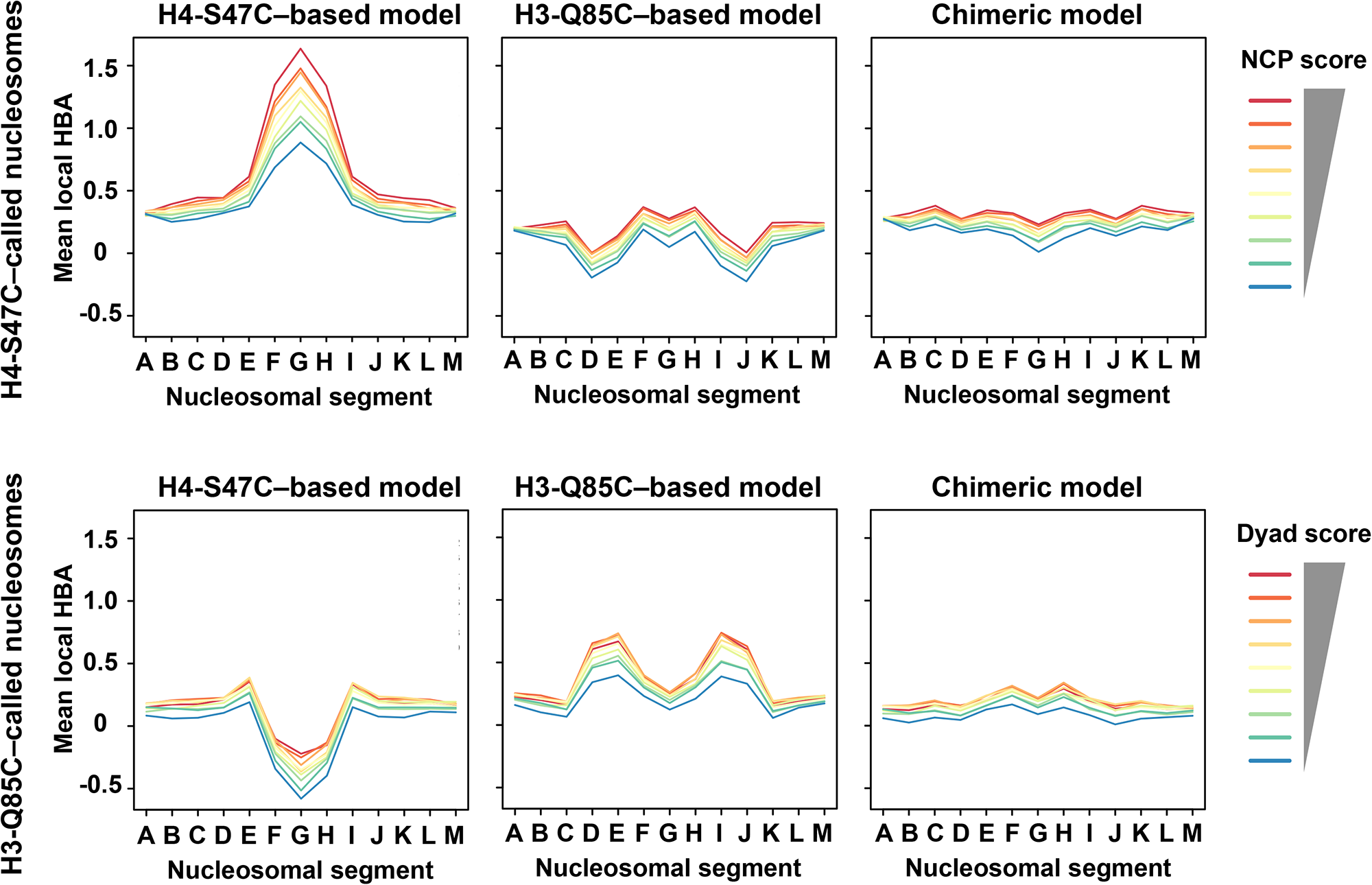
Local HBA analysis of chemically called nucleosomes with distinct statistical models. Nucleosomes called by the methods indicated to the left were grouped according to their detection frequency (NCP score for H4-S47C–called nucleosomes and dyad score for H3-Q85C–called nucleosomes). For each group of nucleosomes, local HBA scores were calculated with the statistical models indicated to the top. Mean local HBA scores for each group were plotted.

H4-S47C–based local HBA scores at the central segments (F, G, and H). In contrast, for the same nucleosomes, H3-Q85C–based local HBA scores were generally low around the sites of H3-Q85C–mediated cleavage (segments D, E, I, and J). Conceptually the same pattern was observed for H3-Q85C–called nucleosomes: H3-Q85C–based local HBA scores were high around H3-Q85C–mediated cleavage sites, whereas H4-S47C–based local HBA scores were low around the dyad. Therefore, each chemical mapping method has sequence biases around cleavage sites and has a tendency to call preferred nucleosomes, which can be located at different translational positions from those preferably called by the other method (**Supplementary Figure S4**).

### Construction of a chimeric model

Consistent with the above results, analysis of nucleosomal DNA sequences clearly showed that each chemical mapping method has specific dinucleotide patterns around cleavage sites in called nucleosomes (**Figures 3A and 3B**, left and center), as previously reported (21). This indicates that prediction of nucleosome positioning with chemical map–based statistical models is affected by these unavoidable biases. To overcome this shortcoming of chemical map–based prediction, we generated a set of chimeric nucleosomal sequences from which we created an ideal nucleosome model (**Figures 3A** and **3B**, right). In order to avoid H4-S47C–associated biases, the central 33 bp segments were derived from the H3-Q85C–called nucleosomes, whereas the flanking 57 bp regions were derived from the H4-S47C–called nucleosomes that are likely to lack H3-Q85C–associated biases. As expected, the biased dinucleotide patterns were not detected in the chimeric nucleosome sequences (**Figure 3B**, right). As expected, method-specific biases in assessing nucleosomal DNA segments were greatly reduced when the chimeric sequence–based statistical model was utilized for local HBA calculations (**Figure 2**, right).

**Figure 3.**
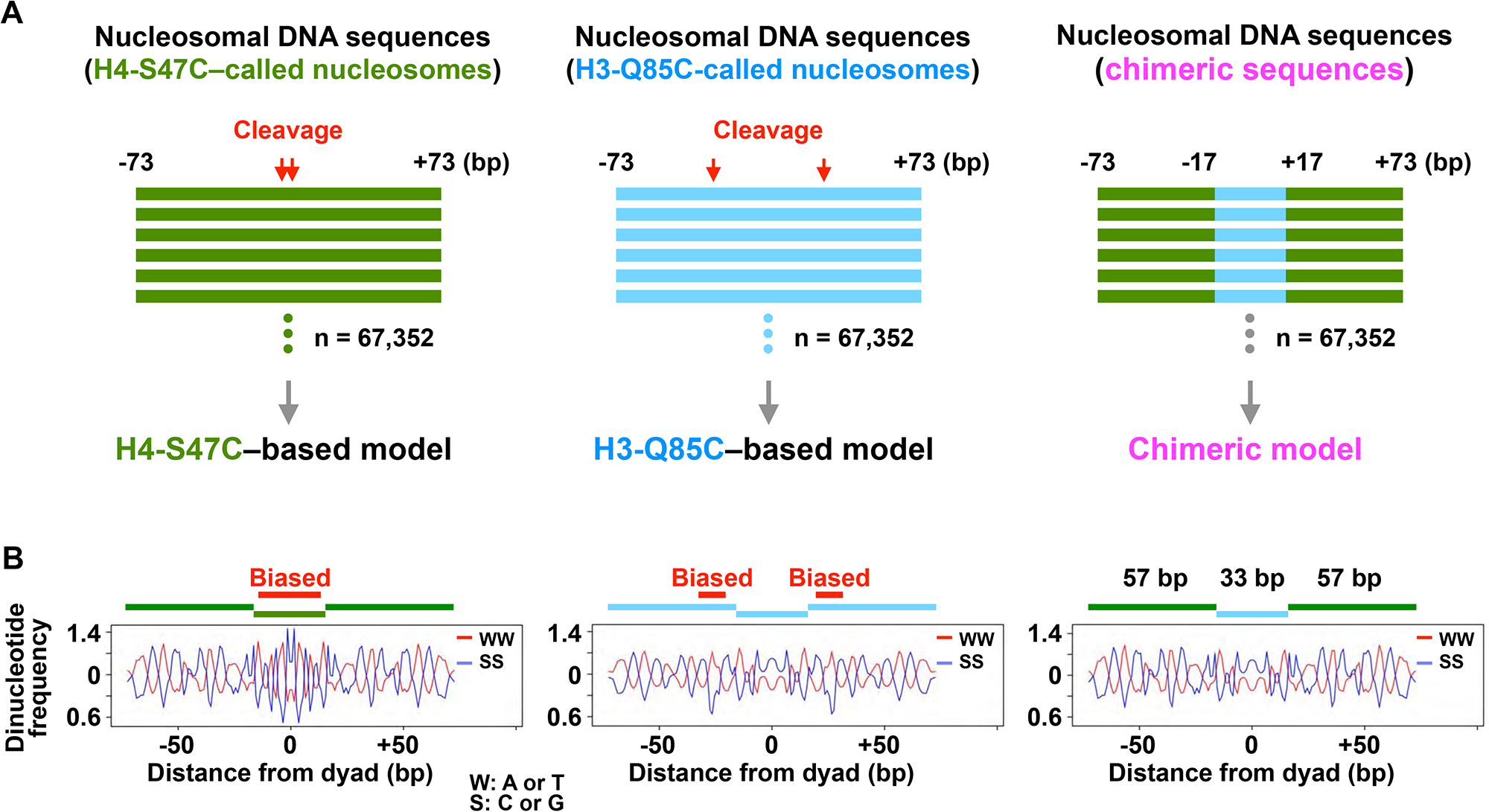
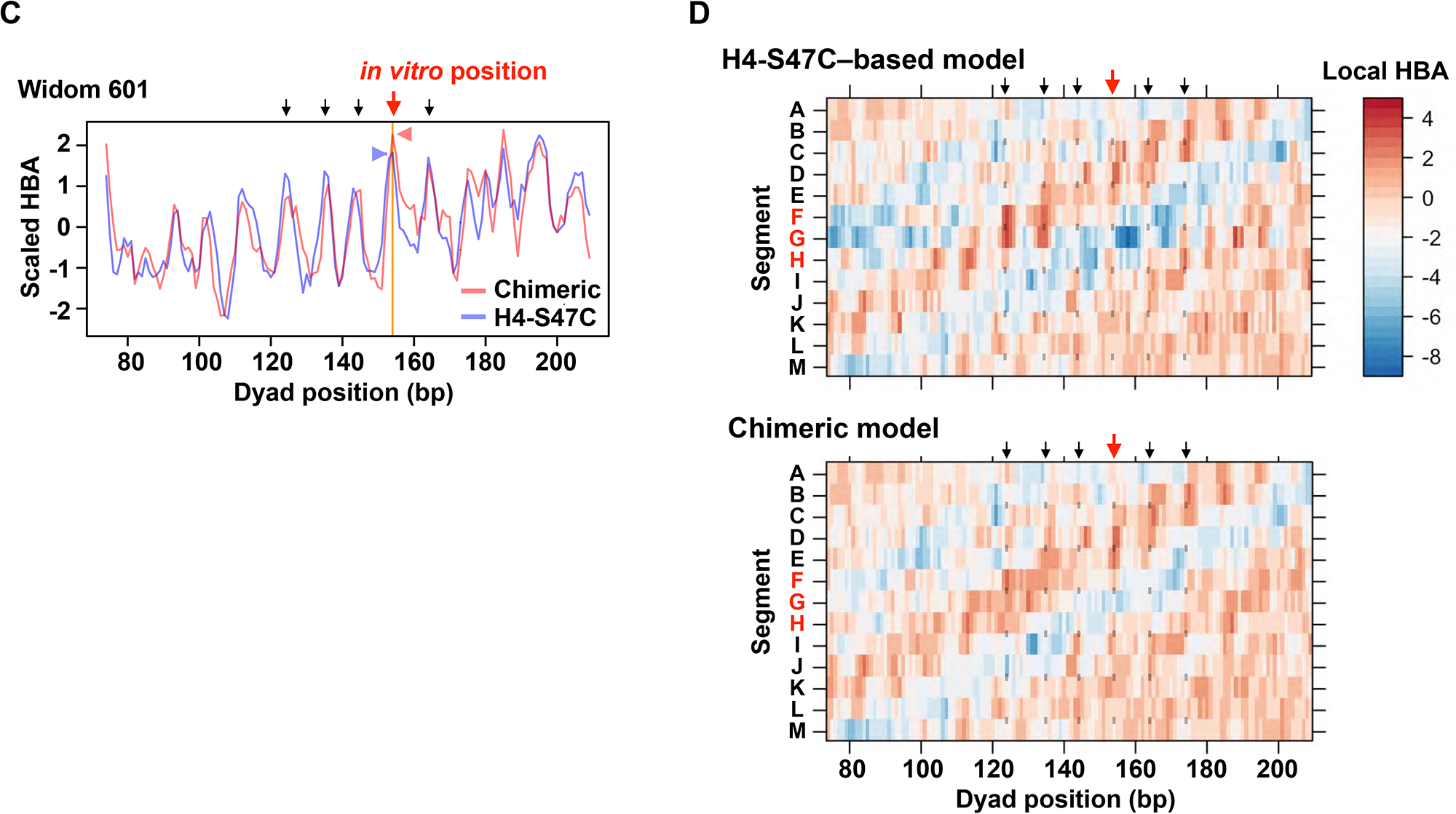
Construction and validation of a chimeric model. (**A**) A scheme for generation of chimeric sequences. From the same number of H4-S47C–called (green) and H3-Q85C–called (light blue) nucleosomal DNA sequences, the outer 57-bp and inner 33-bp sequences were combined to generate chimeric sequences. (**B**) Dinucleotide patterns in nucleosomal DNA sequences. WW refers to the combined frequency of AA, AT, TA, and TT; SS refers to CC, CG, GC, and GG. Intranucleosomal regions with method-specific biases are shown. (**C**) HBA scores along the 282-bp Widom 601 sequence. HBA scores for given 147-bp sequences calculated with the indicated models were assigned to their center positions. HBA scores were scaled to have the same mean and standard deviation for each model. Peak height of HBA scores at the dyad position of the reconstituted nucleosome (red arrow) are indicated with arrow heads. Black arrows indicate potential dyad positions of translationally shifted nucleosomes. (**D**) Local HBA scores along the same sequence as in **C** were calculated with the indicated models and heatmapped at the center position of the tested 147-bp sequences. Red and black arrows and gray points indicate dyad positions as in **C**.

In order to validate the usefulness of the chimeric model, we calculated whole nucleosomal HBA scores along the 282-bp Widom sequence, on which a nucleosome is formed at a specific position *in vitro* (32). Our recent study demonstrated that the translational position of a reconstituted nucleosome on this sequence can be predicted with the previous H4-S47C–based model derived from 67,538 nucleosomes (26). With an interval of ∼10 bp, the model also suggested other potential translational positions with relatively high HBA scores, which could be lowered by improvement of prediction accuracy. Here, we utilized H4-S47C–based and chimeric sequence–based models derived from the same number of nucleosomes (n = 67,352, **Figure 3**) for HBA calculations. The chimeric model–based HBA score at the *in vitro* translational position surpassed that of the matched H4-S47C–based model (**Figure 3C**, red arrow), with lower scores at the other potential positions (black arrows). Local HBA analysis on the same sequence clearly showed that the H4-S47C–based model had a strong preference at the F, G, and H segments, which was reduced by the chimeric model (**Figure 3D**). The improvement in prediction of translational positioning and the reduction in cleavage-related biases rationalize the use of this chimeric model in the assessment of genomic DNA sequences for their suitability for nucleosome formation.

### Asymmetric HBA in protein-coding regions

Using the chimeric model, we first calculated whole nucleosomal HBA scores for 147-bp sequences (**Figure 4A**, upper panel) along yeast protein-coding genes (**Figure 4B**, upper panel). The HBA scores were assigned to the center of the tested 147-bp sequences. The mean HBA scores were then smoothed using a 51-bp centered moving average. Consistent with the previous H4-S47C–based model (26), the chimeric model–based unsmoothed HBA marked highest at the position of Chereji’s +1 nucleosome (21) to which each gene was centered. Unsmoothed mean HBA scores exhibited dramatic oscillations over ∼10 bp intervals, suggesting that the DNA sequence contributes to rotational setting (26). Interestingly, smoothed mean scores (thick green line) peaked slightly upstream of the peak positions of *in vivo* nucleosome arrays, which is consistent with a recent report by Singh et al. (9). Besides the involvement of the remodelers that they studied (9), we hypothesized that the DNA sequence itself may also play unknown roles in nucleosome positioning in transcribed regions.

**Figure 4.**
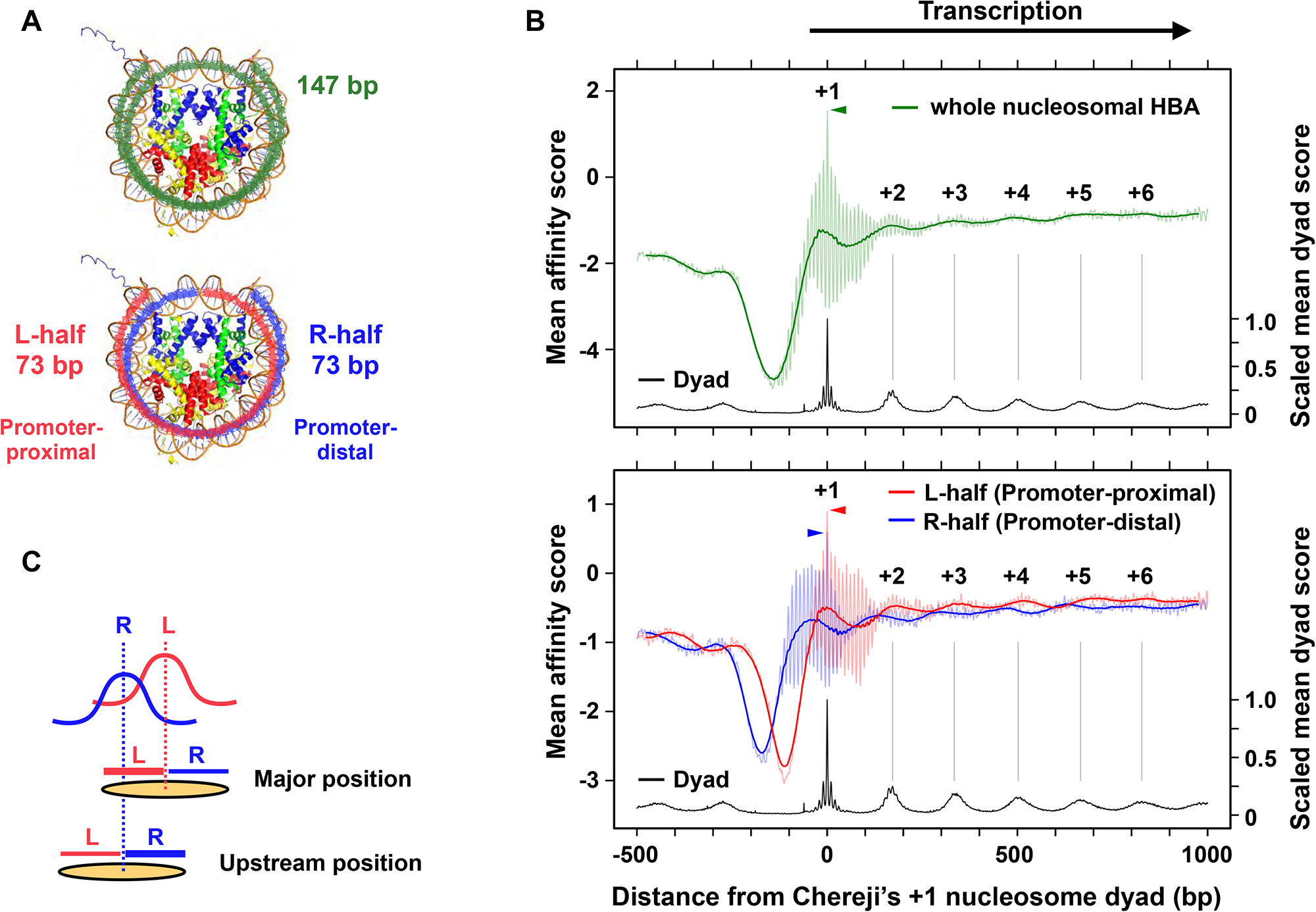
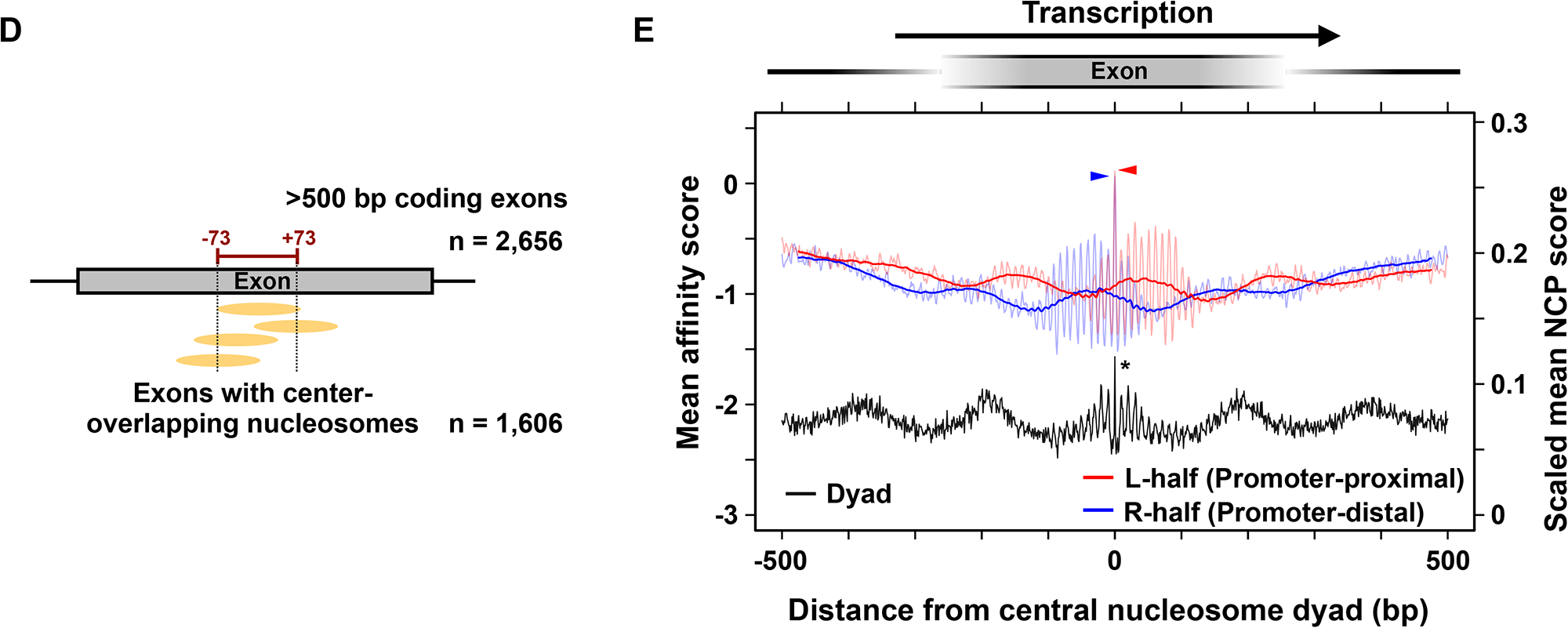
Asymmetric patterns of nucleosome positioning sequences in protein-coding regions. (**A**) Segments of nucleosomal DNA used for HBA calculation. The entire 147-bp nucleosomal sequence (green) and the 73-bp segments for the promoter-proximal (L-half, magenta) and -distal (R-half, blue) halves are indicated on the nucleosome structure (1AOI). (**B**) Mean affinity scores and dyad scores of H3-Q85C–called nucleosomes along yeast protein-coding genes with Chereji’s +1 nucleosome (n = 2,148, > 1.5 kp) (21). HBA scores were assigned to the dyad position of tested sequences (colored thin lines) and smoothed as 51 bp centered moving average (colored thick lines). Approximate peak positions of dyad counts corresponding to +2 to +6 nucleosomes are indicated as vertical gray bars. Height of affinity peaks at the +1 nucleosome position are indicated with arrowheads. (**C**) Interpretation of the shift in affinity peaks for the R-half segment toward the upstream direction. (**D**) Selection of mouse protein-coding exons. > 500 bp RefSeq Select exons with a H4-S47C–called representative nucleosome dyad in the central 147 bp region were selected for HBA analysis. (**E**) Mean affinity scores and NCP scores of H4-S47C–called nucleosomes along murine protein-coding exons are plotted as in the lower panel of **B**. *Scaled mean NCP score at the position of the central nucleosome, which scored 1.0, is intentionally trimmed to clearly show the L-half and R-half scores.

Shifting of the affinity peaks toward the promoter implied that the native sequences have an asymmetric feature in terms of nucleosome formation. To examine this possibility, we split the 147-bp region into two 73 bp segments (designated L- and R-halves) excluding the dyad base pair and calculated chimeric sequence–based HBA scores for each segment (**Figures 1A** and **4A**, lower panel). Unsmoothed mean affinity scores for both halves marked highest at the +1 nucleosome position (**Figure 4B**, lower panel). If nucleosomes do not have a remarkable asymmetric feature, peak positions of smoothed mean HBA scores for both halves would be equivalent. However, the HBA scores oscillated independently along the protein-coding regions. The promoter-proximal half (L-half) generally had higher HBA scores than the downstream half (R-half). Strikingly, affinity peaks for the promoter-proximal half matched well with the positions of the first few nucleosomes, with relatively higher affinity scores downstream. In contrast, affinity scores for the promoter distal segment peaked upstream where parts of the L-half segments of the major nucleosomes were subjected to R-half affinity calculation as depicted in **Figure 4C**. Similar asymmetric oscillation patterns were observed in mouse coding sequences with higher affinity for the promoter-proximal segments (**Figures 4D** and **4E**), implying an underlying fundamental mechanism. Given these results, coding regions are predicted to have higher affinity sequences for the promoter-proximal half of the nucleosome in a periodic manner.

### Impact of complementary dinucleotides on detection of in vivo nucleosomes

To understand the basis of asymmetricity in nucleosome formation, we decided to analyze the relationship between unambiguous short DNA sequences and nucleosome detection frequency. As the nucleosome is a pseudo two-fold symmetric structure (1,2), we assumed that pairs of short nucleotide tracks that are not self-complementary, but complementary to each other (e.g., dinucleotides containing only purines or pyrimidines such as AG/CT), differentially affect the formation or stability of nucleosomes when they are located at the same position. To assess this hypothesis, we selected chemically mapped *in vivo* nucleosomes with particular dinucleotides at particular positions on the top strand (e.g., 5’-AA-3’ at the nucleotide position -47 to -46 relative to the dyad) and those with the complementary dinucleotides at the same position on the same strand (e.g., 5’-TT-3’ at position -47 to -46) (**Supplementary Figure S5**). We then compared these nucleosomes in terms of their detection frequency in chemical cleavage methods.

Although the analyses involving the whole genomic nucleosomes only showed a moderate tendency toward exhibiting the expected asymmetric dinucleotide preference, a clear trend was observed for the nucleosomes located in intergenic regions (**Figure 5** and **Supplementary Figure S6**; for statistics, see **Supplementary Tables S3** to **S6**). This tendency was reasonable because the asymmetric nucleosomes in the protein-coding regions were omitted to extract intergenic nucleosomes. The dinucleotide pair with the most prominent effect was AA/TT (**Figure 5**). The detection frequency of nucleosomes containing AA at specific positions in the left half was generally higher (∼1.05 fold at each position, p < 0.05 in Welch t-test) than of those containing TT at the same positions; the converse was observed in the right half (**Figure 5**, bottom panels). To a lesser extent, similar trends were observed for AG/CT, GA/TC, and GG/CC pairs (**Supplementary Figure S6**). Combined analysis for RR/YY was in agreement with the above results (**Supplementary Figure S7**). These data suggest that RR and YY dinucleotides (typified by AA and TT) in the left and right halves, respectively, of the top strand are favored for nucleosome formation *in vivo*.

**Figure 5.**
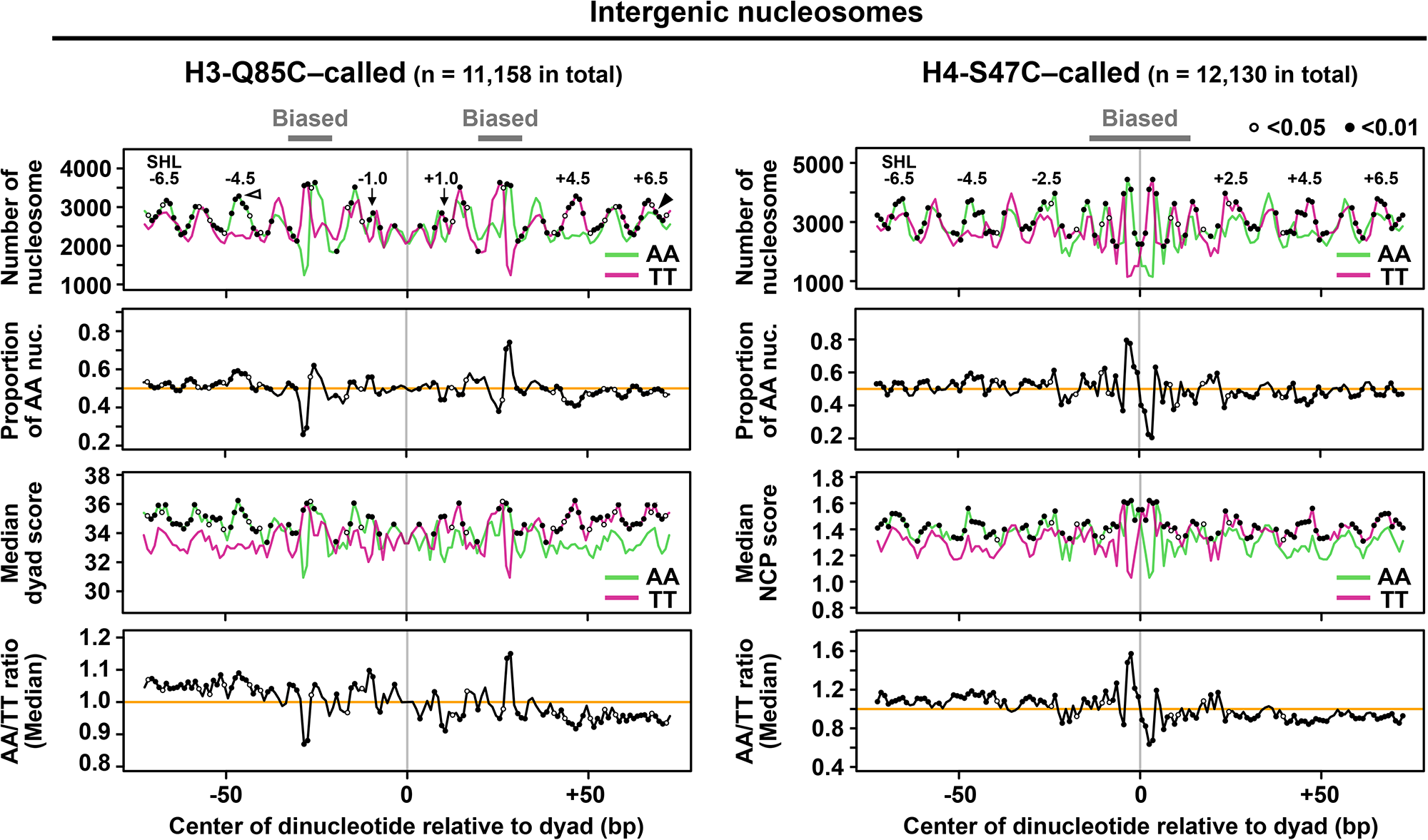
Relationship between AA/TT dinucleotides and nucleosome detection frequency. Analyses were performed on H3-Q85C–called (left) and H4-S47C–called (right) intergenic nucleosomes. The number of selected nucleosomes with AA or TT at each position on the nucleosomal coordinate are shown in the topmost panels. White and black arrow heads indicate the dinucleotide positions exemplified in **Supplementary Figure S5**. Regions with method-specific biases and SHLs of interest are indicated. The proportion of nucleosomes containing AA at distinct positions in the selected nucleosomes with AA or TT at the same position are shown in the second panels. Median detection frequency of the selected nucleosomes with AA or TT at distinct positions are shown in the third panels. Significant differences (p < 0.05 or p < 0.01, Welch t-test) in each comparison are indicated as open and closed circles, respectively, in all the relevant plots. The ratio of median detection frequency (AA- versus TT-containing nucleosomes) are plotted in the bottom panels. For whole genomic nucleosomes and other dinucleotide pairs, see **Supplementary** Figures S6 and S7.

The general, non-periodic preference for RR/YY dinucleotides in each side of the nucleosome was unexpected because the frequency of these dinucleotides (at least for the typical AA/TT pair) was apparently periodic along the nucleosomal coordinate. To elucidate this non-periodic preference, we analyzed nucleosomal DNA by ranking the sixteen kinds of dinucleotide with respect to their proportion at each nucleotide position (**Figure 6** and **Supplementary Figure S8**). By considering proportion landscapes of dinucleotide rankings, rather than just examining frequencies of each dinucleotide (11,21), we are able to focus on intranucleosomal regions where limited patterns of sequences are selectively required. In addition, mapping of particular dinucleotides on the proportion landscapes allows us to examine how they are distributed with respect to the nucleosomal coordinate.

**Figure 6.**
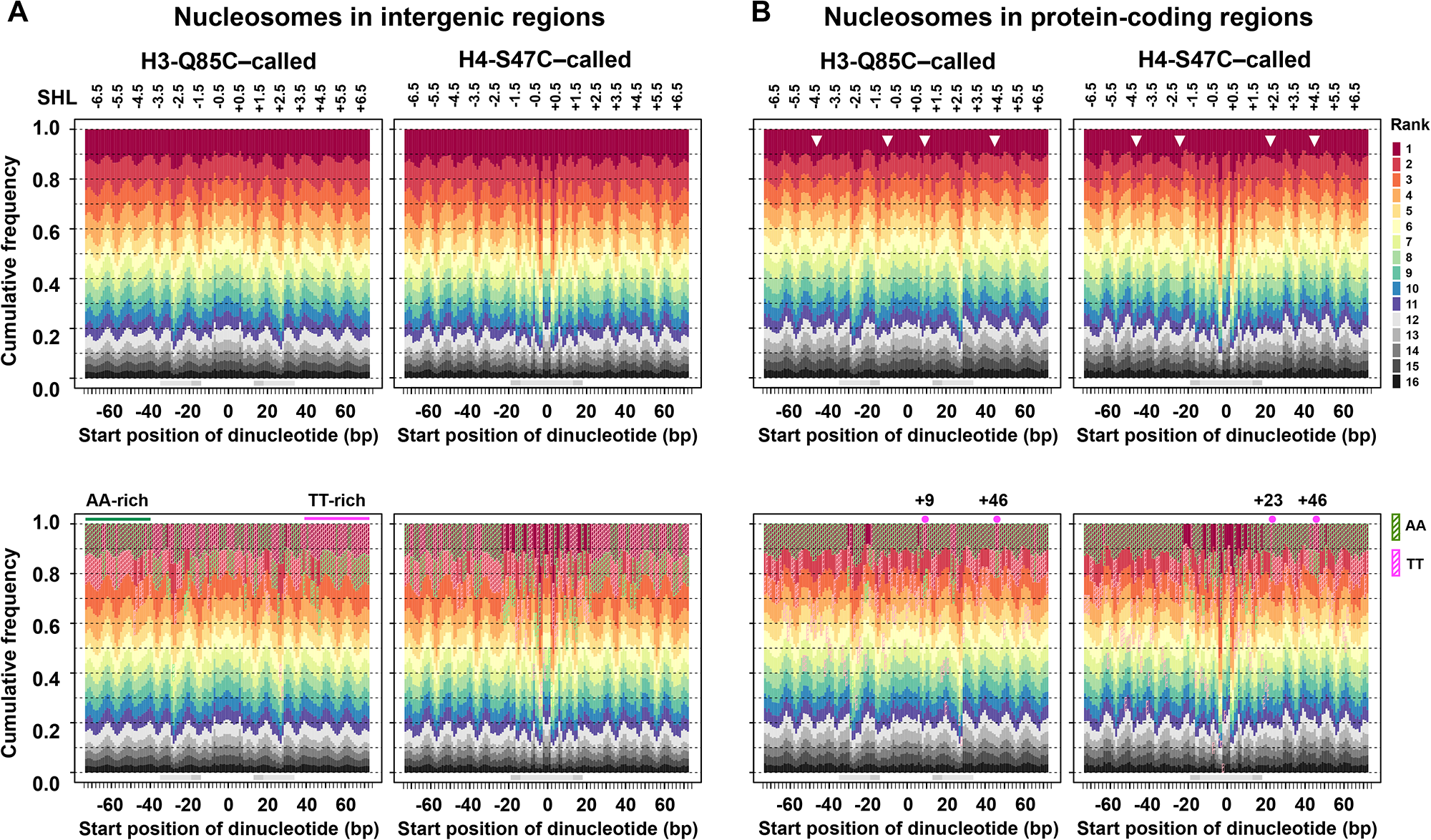
Visualization of dinucleotide ranking along the nucleosomal DNA coordinate. Analyses were performed on chemically detected nucleosomes locating in intergenic (**A**) and protein-coding (**B**) regions. Color scales for **A** and **B** are shown on the right side of **B**. Dinucleotide ranks at each starting position are colored to draw proportion landscapes (top panels). Nucleosomal regions with the cleavage biases (light gray) and those potentially affected by chimera generation (dark gray) are indicated. The location of AA (green) and TT (magenta) are indicated in the proportion landscapes shown in the bottom panels. For the intergenic nucleosomes (**A**), AA-rich and TT-rich regions are indicated. For the protein-coding nucleosomes (**B**), results of +2 nucleosomes are shown. For +1, +3, and +4 nucleosomes, see **Supplementary Figure S8**. Arrow heads in the upper panels indicate the -47, -24, -10, +9, +23, and +46 nucleotide positions. In the lower panels, the +9, +23, and +46 nucleotide positions, where AA does not rank first, are indicated. Positions -24 and +23 for H3-Q85C–called nucleosomes and -10 and +10 for H4-S47C–called nucleosomes are not indicated to avoid futile examination around cleavage sites.

In nucleosomes within the intergenic regions, in which transcription or translation is not relevant, selection of dinucleotides appeared to be limited at histone contact sites (e.g., SHL -6.5) where the sum proportion of the top five dinucleotides accounted for over 50% of the nucleosomes analyzed (**Figure 6A**, upper panels). In contrast, freedom of dinucleotide selection appeared higher in the other intranucleosomal regions (non-histone contact sites). The dinucleotide proportion landscapes and mapping of AA and TT (**Figure 6A**, lower panels) showed that the H4-S47C–mediated nucleosome calling method (right) has greater sequence biases around cleavage sites than the H3-Q85C–mediated calling method (21). This suggests that the H4-S47C–mediated method is more likely to call nucleosomes that are located in minor translational positions but are suitable for cleavage. Consistent with the observed correlation of AA/TT dinucleotides with nucleosome detection frequency (**Figure 5**), AA and TT often ranked first in the left and right sides of intergenic nucleosomes, respectively, especially in the H3-Q85C–called nucleosomes (**Figure 6A**, lower panels). This tendency was even observed at the non-histone contact sites. Therefore, these dinucleotides can be accommodated throughout the nucleosome and should not be recognized as unpreferred at non-histone contact sites.

For nucleosomes located in coding sequences (**Figure 6B**, upper panels), six kinds of dinucleotide were required to constitute a 50% proportion at histone contact regions, suggesting the influence of codons on the selection of DNA sequences for DNA wrapping. In addition, frequency landscapes of first ranked dinucleotides at SHL ± 1.0, ± 2.5, and ± 4.5 were different on the left and right (white arrows). Note that SHL ± 1.0 and ± 2.5 can only be evaluated in the H3-Q85C–called and H4-S47C–called nucleosomes, respectively, due to the cleavage biases (gray horizontal bars). The lower proportion of the first dinucleotides in the right side suggested a compromise in DNA sequence selection in the promoter-distal half. We noticed that the overall frequency of AA in the top strand (sense strand) of wrapped DNA was ∼1.6-fold higher than that of TT, and AA ranked first at almost all nucleotide positions on the nucleosomal coordinate, including the right (promoter-distal) side (**Figure 6B**, lower panels). Consistent with its lower overall frequency, TT ranked second to eighth even in the unbiased regions. The highest proportion of AA in the promoter-distal half may be associated with the lower affinity of DNA on this side, as suggested by the detection frequency analysis of intergenic nucleosomes (**Figure 5**).

## DISCUSSION

By analyzing cysteine-mediated cleavage and local DNA sequences of chemically called nucleosomes with statistical models, we demonstrated that currently available chemical mapping methods have specific sequence preferences for cleavage and thus tend to call nucleosomes in a biased manner. In these methods, the detection frequency of a nucleosome located at a specific position may be determined by the balance between the abundance of nucleosomes and cleavage efficiency. A major determinant of chemical cleavage efficiency was found to be the local DNA sequence around the cleavage sites. Additionally, all other local segments of the nucleosomal DNA had a minor but significant effect on cleavage. This observation suggests that the abundance or stability of a nucleosome is at least in part determined by the overall suitability of wrapped DNA for nucleosome formation, as previously mentioned (33). Given the above findings, we created a less-biased statistical model from chimeric nucleosomal DNA sequences with the intention of studying the roles of DNA sequences in nucleosome positioning *in vivo*.

Analysis with the chimeric model revealed that in yeast protein-coding sequences there is a periodic occurrence of 73 bp DNA sequences that are suitable for wrapping around histones in the promoter-proximal half. The coincidence of affinity peaks for the promoter-proximal half and the positions of the first few nucleosomes in yeast imply that the cells exploit the intrinsic affinity for this half to modulate the positioning of nucleosomes. Note that the favored sequences were not found as single distinct positions, but rather as regions in which nucleosomes can be positioned redundantly. Unsmoothed scores indicated there is ∼10-bp oscillation of affinity that directs rotational setting of redundant positioning. The first few nucleosomes often incorporate histone variant H2A.Z (34), which destabilizes nucleosome positions (35). In addition, these nucleosomes are vulnerable to impairment of histone chaperones (7,10,36,37). Therefore, affinity peaks for the promoter-proximal half matching *in vivo* nucleosome positions could be related to these specific features. In contrast, for the following nucleosomes, high affinity sequences for the promoter-proximal half peaked downstream of *in vivo* nucleosomes, suggesting a difference in regulation.

When transcription occurs, the RNA polymerase II-containing transcription machinery coming from the promoter must first unwrap the nucleosomal DNA in the promoter-proximal half in order to transcribe the encoded information (15,16,38,39). Before pealing the whole nucleosomal DNA off the histones, the promoter-proximal half needs to rewrap around, or at least reestablish contact with, the preexisting histones to avoid their loss during transcription. Indeed, maintenance of preexisting nucleosomes in transcription was experimentally demonstrated *in vivo* (40,41). Moreover, recent cryo-electron microscopy analysis revealed that the rewrapping of DNA first occurs in the promoter-proximal half of the nucleosome (42). Loss of histones is often caused by misfunction of histone chaperones accompanying the polymerase. This could lead to loss of locus-specific posttranslational modifications (7,36) and to activation of cryptic transcription (37,43,44), both of which could perturb epigenetic regulation. Considering the DNA sequence patterns presented here, we propose that arrangement of nucleosome positions in protein-coding regions may also rely on the intrinsic suitability of DNA, especially for the promoter-proximal half of the nucleosome. The similar periodic patterns found in the murine coding sequences strongly suggest that the underlying mechanism is conserved among eukaryotes.

The overall AA-richness in the sense strand of yeast protein-coding regions is consistent with the existence of multiple intragenic sequences with high affinity for the promoter-proximal half. Considering that nucleosomes are often redundantly detected at neighboring, overlapping positions (11,21,22,45), that transcriptional induction is associated with a shift in nucleosome position toward the downstream direction (46–49), and that rewrapping in the wake of RNA polymerase II potentially causes a register shift of wrapped DNA (15,16,42,50), DNA sequences with intrinsic affinity to the promoter-proximal half should be present redundantly, especially in the downstream sequence. Once rewrapping of DNA in the upstream half determines a nucleosome’s translational position, post-transcriptional rewrapping of the other half could occur passively. In this scenario, optimization of the coding sequence for the promoter-proximal half is reasonable for histone maintenance. Maintenance of preexisting histones that are post-translationally optimized for transcription provides the benefit of secure transcription cycles that continue to suppress cryptic transcription. This eukaryote specific requirement may have exerted evolutionary pressure on spontaneous DNA sequence alterations and yielded the asymmetric patterns of nucleosome positioning sequences. Moreover, it would be worth exploring whether nucleosomes are relocated to more favorable home positions after transcription.

Although the affinity scores for the promoter-distal half in the coding region are relatively low compared to those for the proximal half, they are much higher than those in the nucleosome depleted regions located upstream of the genes. This suggests that the lower affinity promoter-distal half may still be suitable for nucleosome formation. At the promoter-distal histone contact regions SHL + 1.0, + 2.5, and + 4.5, AA was not ranked first despite the AA-richness in the sense strand (see nucleotide positions + 9, + 23, and + 46 in **Figure 6B** lower panel). As AA was ranked third or fourth at these positions of intergenic nucleosomes, local sequences containing AA can have a negative impact on nucleosome positioning at these intranucleosomal regions. This in turn suggests that the promoter-distal sequences also contribute to stabilizing nucleosome positions. Near SHL ± 2.5 and ± 4.5, structural deformation of DNA is associated with nucleosome remodeling (19,51,52). In addition, a molecular dynamics simulation study revealed that DNA at SHL ± 1.0 is attached by histone H3 “latch” residues and that its distortion concomitantly occurs with unwrapping of the terminal regions of nucleosomal DNA locating in the other half (19). Collectively, further attempts to understand nucleosome positioning *in vivo* should take local sequences into account.

The impact of AA/TT on nucleosome positioning in yeast has previously been reported by Ioshikhes et al., showing that the first few nucleosomes in the gene could be predicted by the frequency of AA and TT dinucleotides (53). Using nucleosomal detection frequency as an indicator of *in vivo* nucleosome abundance, we found that AA and TT have a role in nucleosome stabilization when these dinucleotides are located in the top strand’s right and left halves, respectively. One interesting point regarding the impact of AA/TT is that their involvement appears to be independent of the periodic DNA-histone contacts. Of the six geometric parameters for the DNA base steps, AA and TT completely differ in tilt angle: AA and TT hardly tilt in the positive and negative angles, respectively (54). Other RR/YY base steps have similar, but less pronounced, characteristics. Thus, we speculate that these dinucleotides contribute to moderation of the physiological structure of wrapped DNA to stabilize nucleosomes *in vivo*.

Although our chimeric approach successfully reduced the prediction biases, the constructed model could still be affected by unnatural local sequences at the chimeric junctions and the unavoidable ambiguity of chemical mapping methods in nucleosome calling. In addition, the currently available chemical mapping methods may not require the target nucleosomes to be fully wrapped, as they cleave DNA in the internal regions (around dyad and SHL ± 2.5) of the nucleosome. Thus, further clarification of the importance of DNA sequences in nucleosome positioning should be undertaken by development of a novel high-resolution mapping method of fully-wrapped nucleosomes, preferably with unbiased intranucleosomal sequences.

## DATA AVAILABILITY

Experimental codes for this study are available online (https://github.com/hkatomed/Katoetal2023).

## SUPPLEMENTARY DATA

Supplementary Data are available at XXX.

Supplementary Tables S1 and S2: Tables_S1_S2.xlsx

Supplementary Table S3: Table_S3.xlsx

Supplementary Table S4: Table_S4.xlsx

Supplementary Table S5: Table_S5.xlsx

Supplementary Table S6: Table_S6.xlsx

## SUPPLEMENTARY FIGURE LEGENDS

**Supplementary Figure S1.** Mean cleavage count along the nucleosomal coordinate for each local HBA group. Lines for lower local HBA groups are overlaid on those for higher score groups. Locations of the segments used for local HBA calculation and nucleosome grouping are indicated as horizontal bars at the bottom. *The results for the D and F segment groupings are shown in **Figure 1D**.

**Supplementary Figure S2.** Cleavage counts at indicated nucleotide positions in each local HBA group. *The results of left side cleavage for the D and F segment groupings are shown in **Figure 1E**.

**Supplementary Figure S3.** Mean cleavage count along the nucleosomal coordinate for selected nucleosomes with highest local HBA at the F or H segments. *The result for the F segment grouping of the nucleosomes with highest local HBA scores at the H segment is shown in **Figure 1H**.

**Supplementary Figure S4.** Biased calling of *in vivo* nucleosomes by chemical mapping methods. Nucleosomes are positioned along the genome redundantly with varying abundance. Dark brown represents nucleosomes at major positions; brighter colors suggest minor positions. An ideal mapping method would be expected to always call nucleosomes locating at major positions with the highest detection frequency. In practice however, if the local sequences are not suitable for chemical cleavage, currently available mapping methods would call minor nucleosomes with preferable sequences around the cleavage sites. Note: only representative nucleosomes are shown here as being called. More minor nucleosomes (redundant nucleosomes) are also detected with method-specific detection frequency.

**Supplementary Figure S5.** Selection of nucleosomes with particular dinucleotides starting at distinct positions and comparison of nucleosome detection frequency. (**A**) For a test nucleosome dataset (e.g., H3-Q85C–called intergenic nucleosomes) with Watson strand sequences and respective nucleosome detection frequency scores, a duplicate dataset with the same detection frequency scores but with reverse complement sequences was produced. The original and duplicate datasets were then combined. Using the combined nucleosome dataset, nucleosomes with particular dinucleotides (e.g., AA) or the complementary dinucleotide (e.g., TT) starting at distinct nucleotide positions (e.g., -47 or +69 in this figure) were selected. (**B**) For the pairs of selected nucleosomes, the Welch t-test was used to examine whether both nucleosome sets have significant differences in nucleosome detection frequency. The comparison was made for all possible dinucleotide starting positions as summarized in **Figure 5** and **Supplementary Figures S6** and **S7**.

**Supplementary Figure S6.** Relationship between dinucleotides and nucleosome detection frequency. Analyses were performed on H3-Q85C–called (left panels) and H4-S47C–called (right panels) nucleosomes as in **Figure 5**, in which results of AA and TT containing intergenic nucleosomes (as marked with asterisks in this figure) are selectively shown. For each chemical mapping method, whole genomic (left) and intergenic (right) nucleosomes were analyzed. Dinucleotide pairs considered for the analysis are indicated (e.g., AA vs. TT).

**Supplementary Figure S7.** Relationship between RR/YY dinucleotides and nucleosome detection frequency. Analyses were performed on H3-Q85C–called (left panels) and H4-S47C–called (right panels) nucleosomes as in **Figure 5** and **Supplementary Figure S6**. Nucleosomes were selected when they have RR (either of AA, AG, GA, or GG) or YY (CC, CT, TC, or TT) dinucleotides at distinct positions.

**Supplementary Figure S8.** Dinucleotide ranking analysis for the nucleosomes in protein-coding regions. Dinucleotide ranking for +1 to +4 nucleosomes is shown as in the right panel of **Figure 6**, in which the results for +2 nucleosomes (*) are presented.

## SUPPLEMENTARY TABLE TITLES

**Supplementary Table S1.** Results of Pearson’s correlation test.

**Supplementary Table S2.** Fold difference of median cleavage counts between highest and lowest local HBA groups.

**Supplementary Tables S3.** Statistics for Figure 5 and Supplementary Figures S6 and S7 (whole genomic H3-Q85C–called nucleosomes).

**Supplementary Tables S4.** Statistics for Figure 5 and Supplementary Figures S6 and S7 (whole genomic H4-S47C–called nucleosomes).

**Supplementary Tables S5.** Statistics for Figure 5 and Supplementary Figures S6 and S7 (intergenic H3-Q85C–called nucleosomes).

**Supplementary Tables S6.** Statistics for Figure 5 and Supplementary Figures S6 and S7 (intergenic H4-S47C–called nucleosomes).

