## Supplemental figures for "Asymmetric patterns of nucleosome positioning sequences in protein-coding regions"

### Figure S1

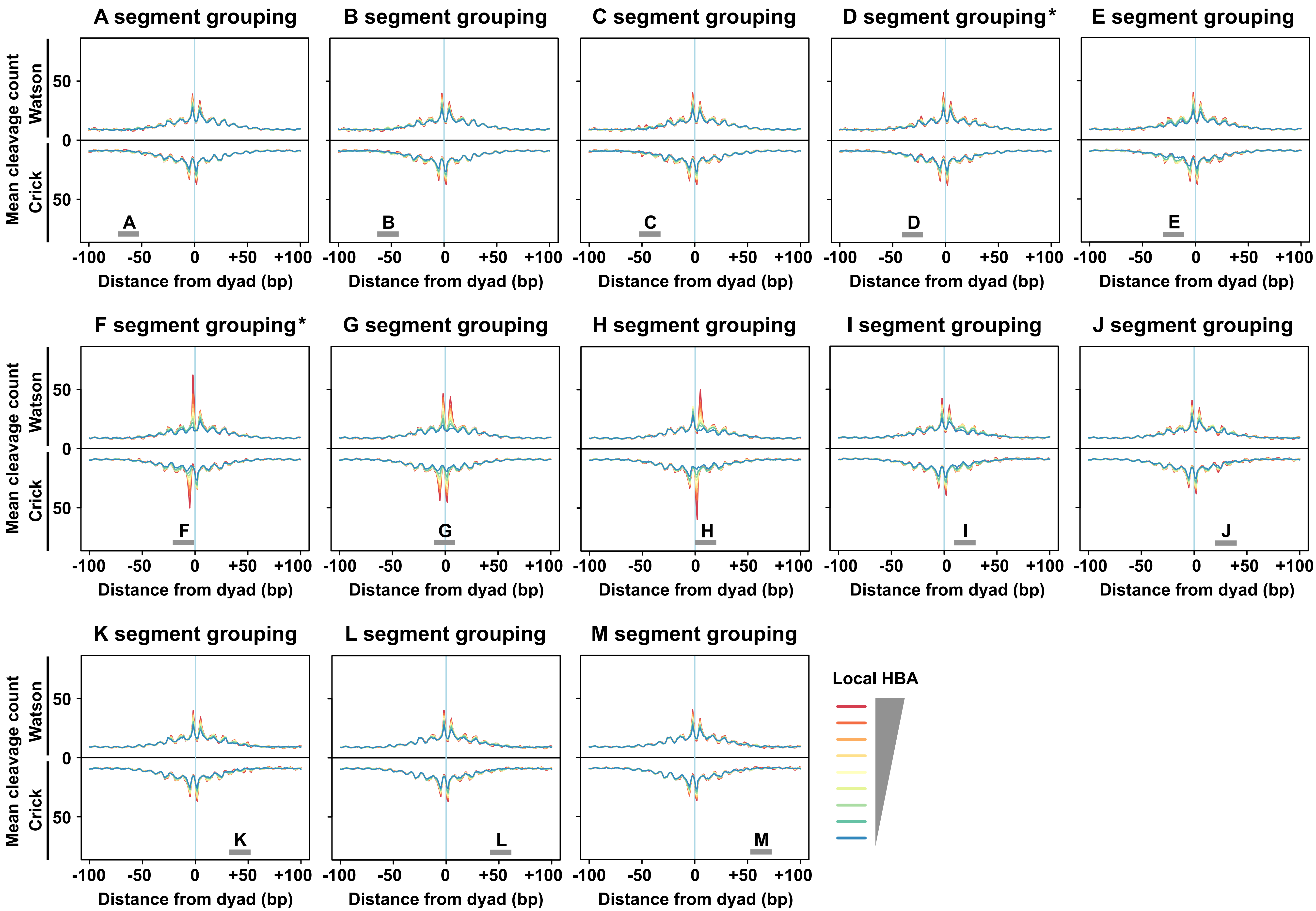
A segment grouping
B segment grouping
C segment grouping
D segment grouping*
E segment grouping
F segment grouping*
G segment grouping
H segment grouping
I segment grouping
J segment grouping
K segment grouping
L segment grouping
M segment grouping
Mean cleavage count
Watson
Crick
50
0
50
-100
-50
0
+50
+100
Distance from dyad (bp)
A
B
C
D
E
F
G
H
I
J
K
L
M
Local HBA

### Figure S2

#### Crick strand (-5)

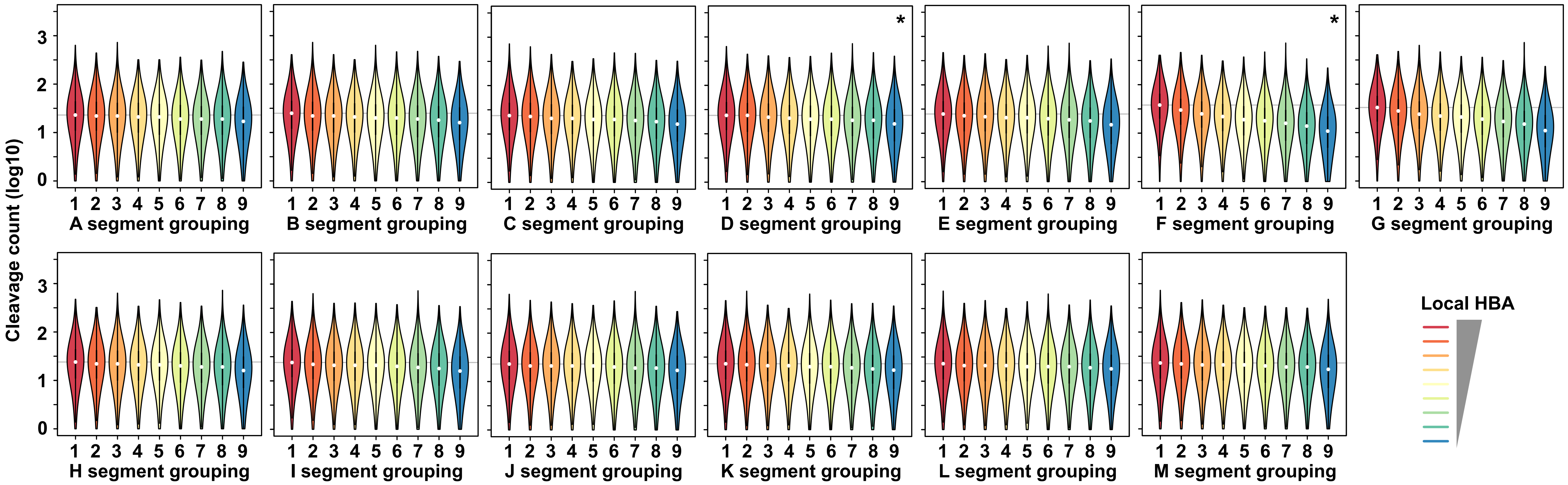

#### Watson strand (-2)

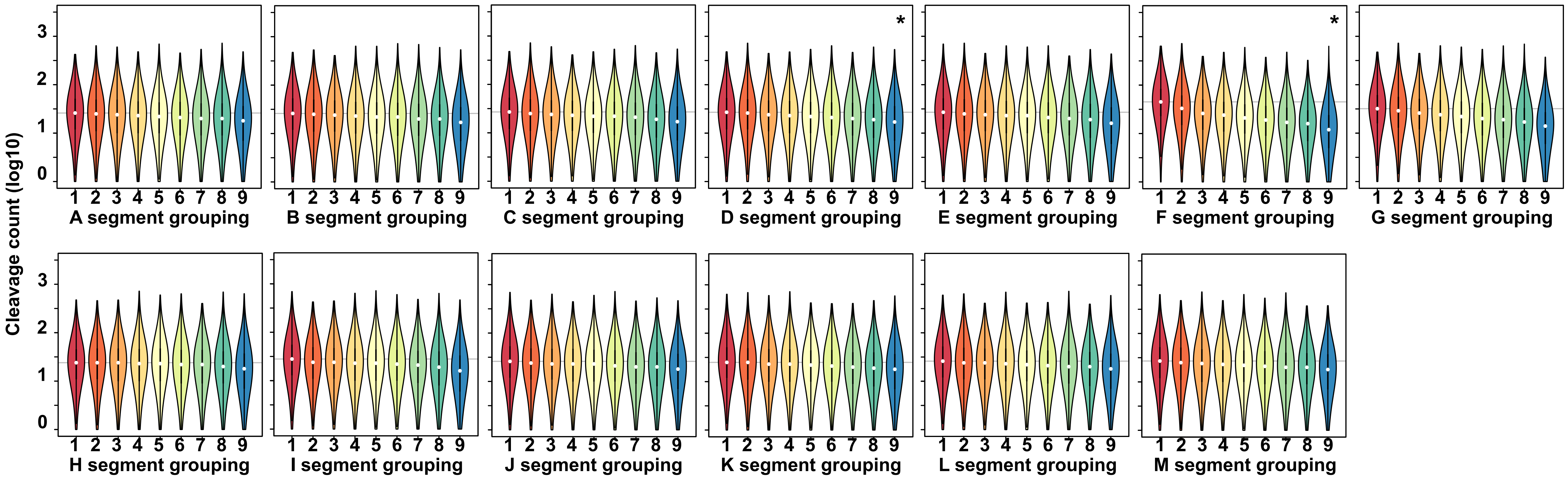

### Figure S2, continued

#### Crick strand (+2)

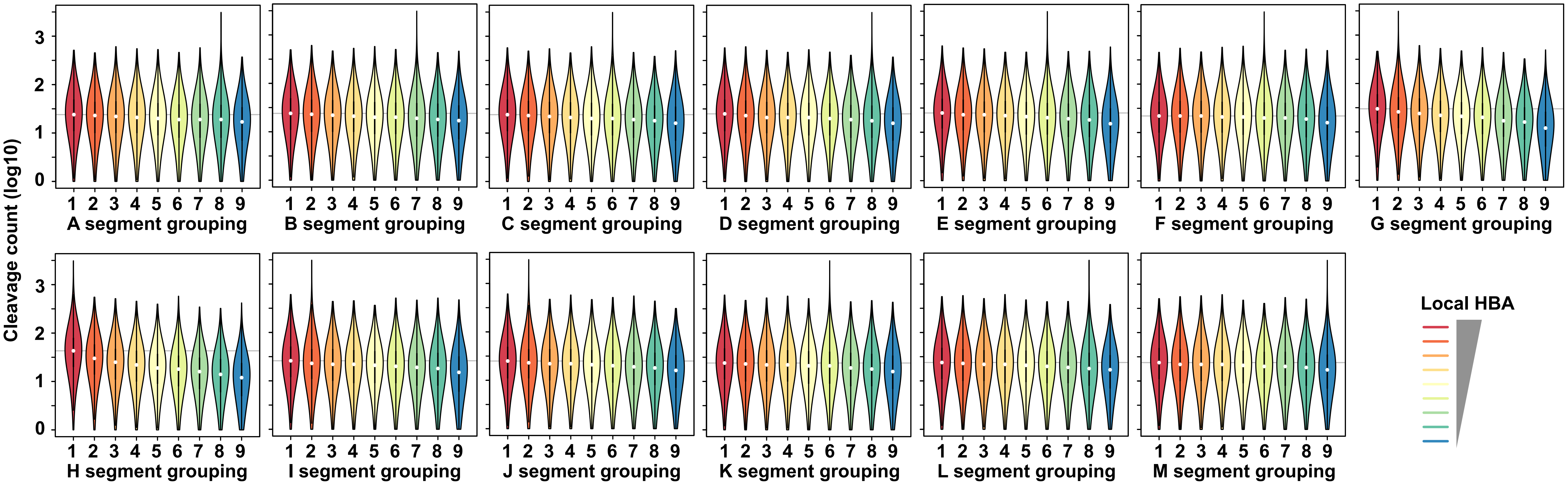

#### Watson strand (+5)

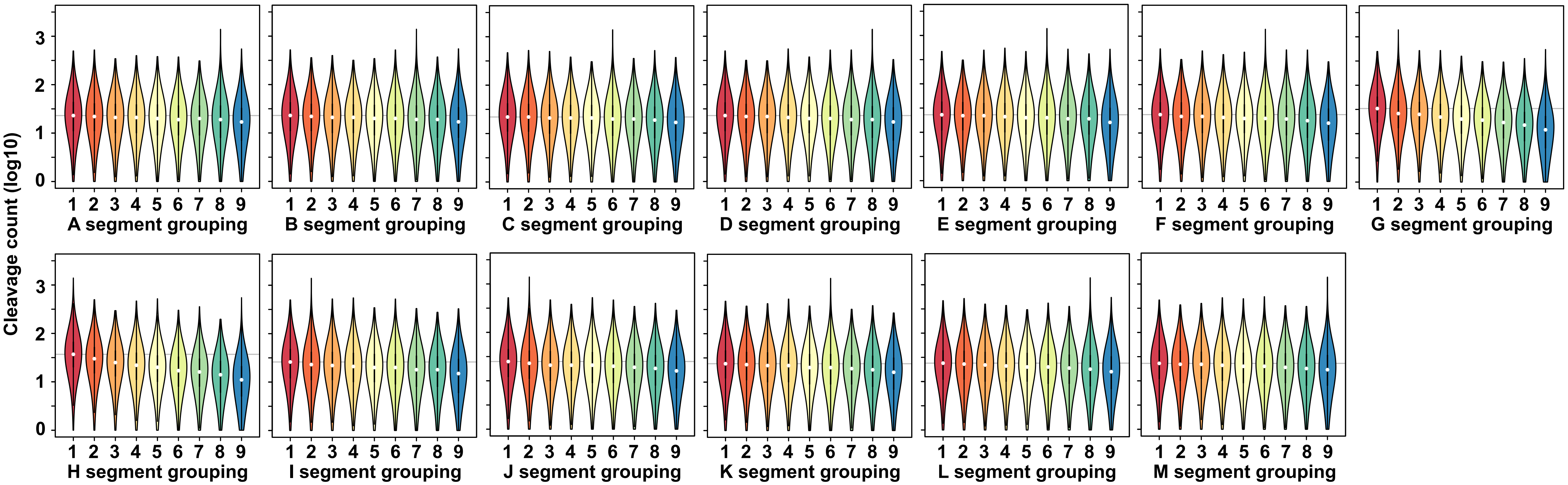

**Figure S3**

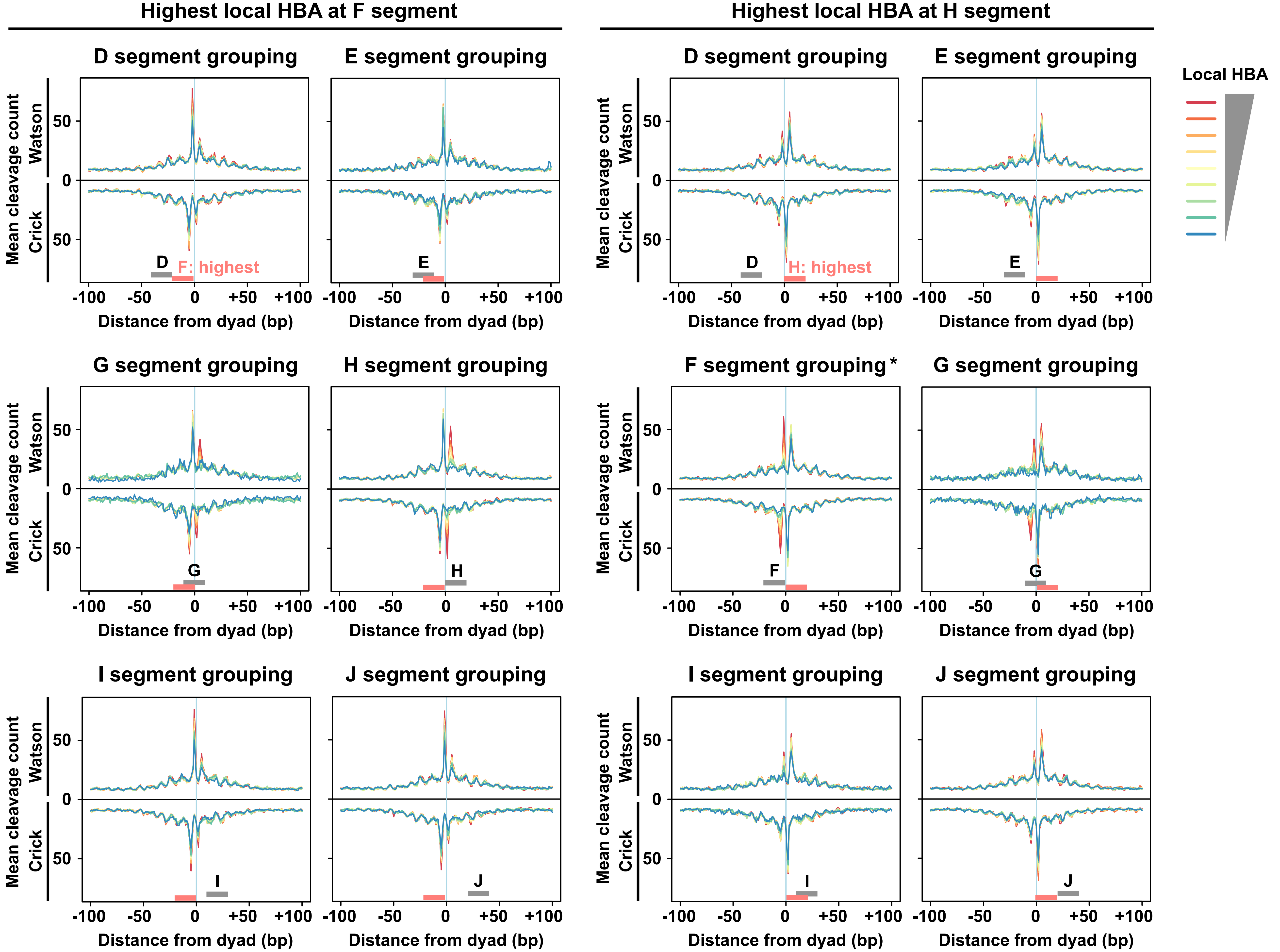
Highest local HBA at F segment
Highest local HBA at H segment
D segment grouping
E segment grouping
G segment grouping
H segment grouping
I segment grouping
J segment grouping
F segment grouping *
Local HBA
Mean cleavage count
Watson
Crick
50
0
D
E
F
G
H
I
J
F: highest
H: highest
-100
-50
+50
+100
Distance from dyad (bp)

Figure S4

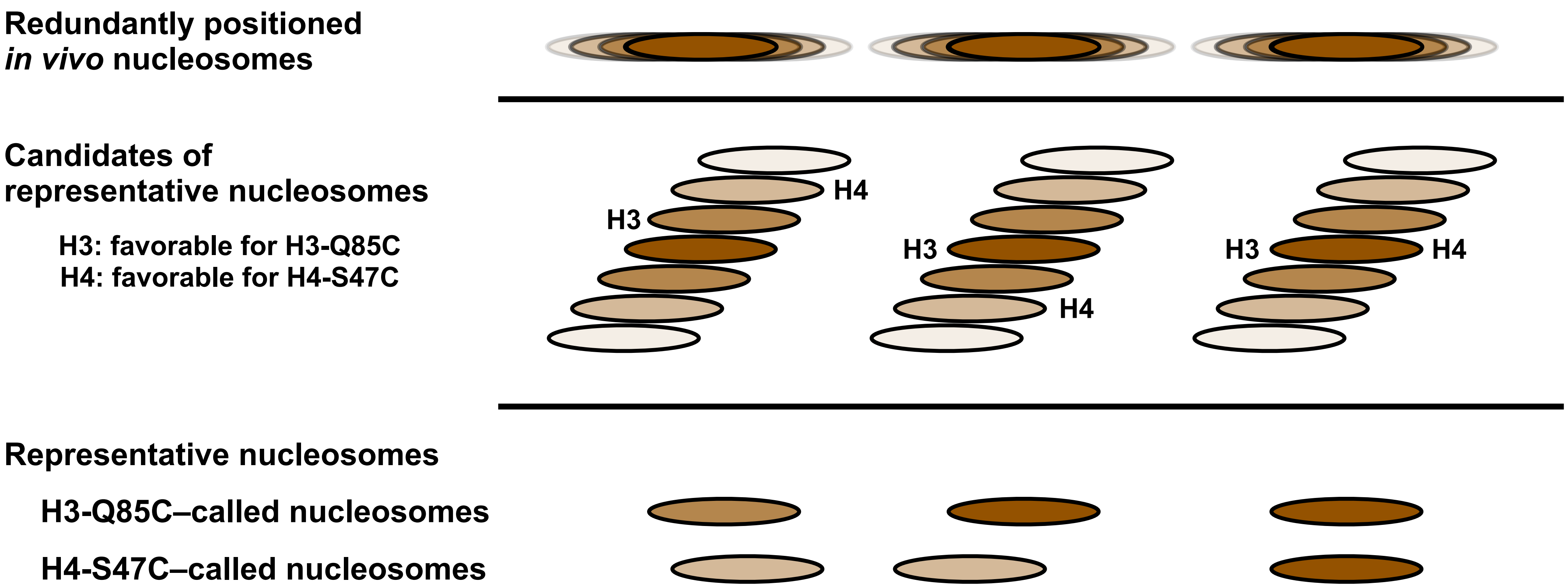
Redundantly positioned
*in vivo* nucleosomes
Candidates of
representative nucleosomes
H3: favorable for H3-Q85C
H4: favorable for H4-S47C
H3
H4
H3
H4
H3
H4
Representative nucleosomes
H3-Q85C–called nucleosomes
H4-S47C–called nucleosomes

Figure S5

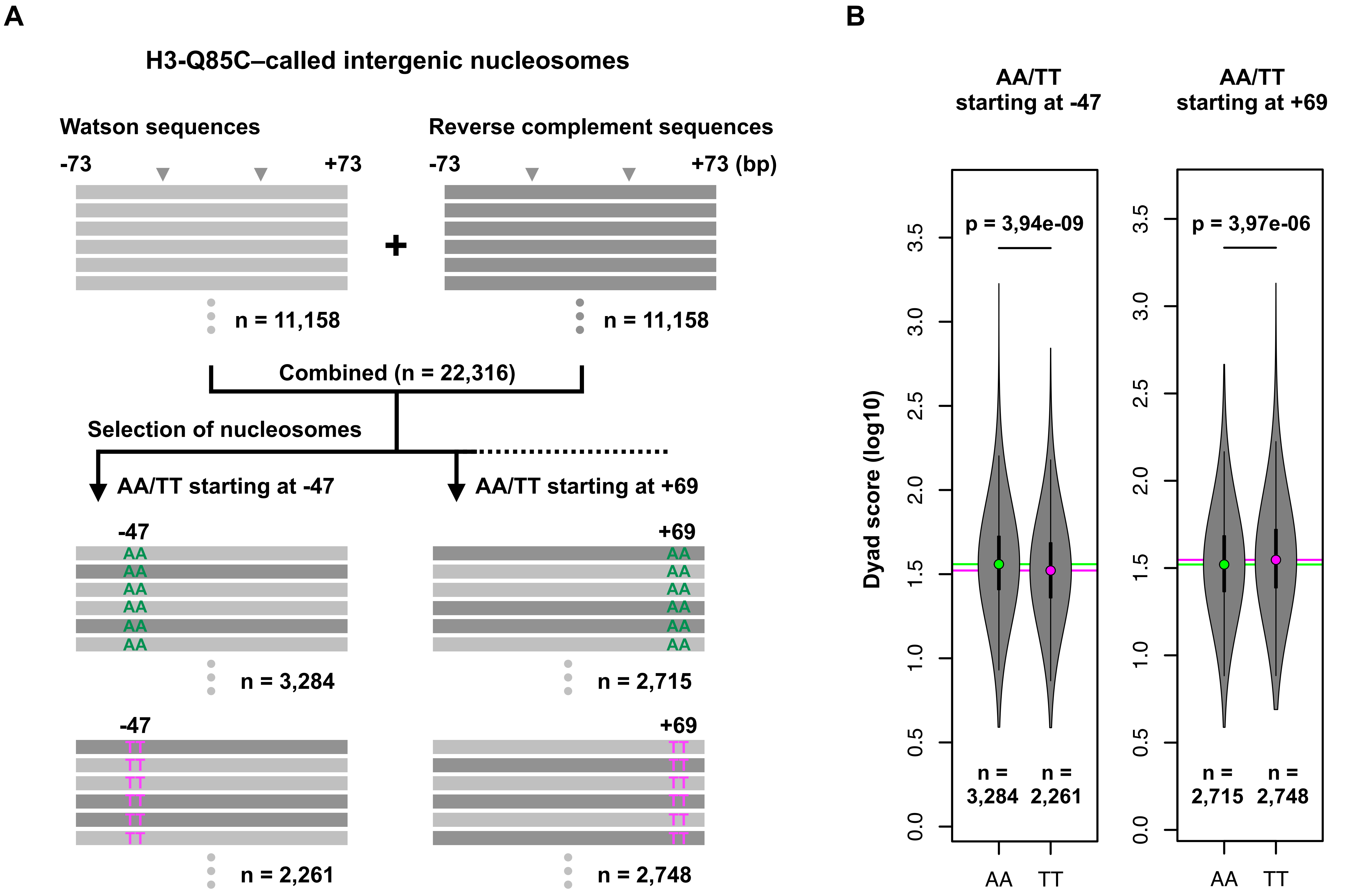
A
H3-Q85C–called intergenic nucleosomes
Watson sequences
Reverse complement sequences
-73
+73
-73
+73 (bp)
+
n = 11,158
n = 11,158
Combined (n = 22,316)
Selection of nucleosomes
AA/TT starting at -47
AA/TT starting at +69
-47
+69
AA
n = 3,284
n = 2,715
-47
+69
TT
n = 2,261
n = 2,748
B
AA/TT starting at -47
AA/TT starting at +69
p = 3,94e-09
p = 3,97e-06
Dyad score (log10)
0.0
0.5
1.0
1.5
2.0
2.5
3.0
3.5
n = 3,284
n = 2,261
n = 2,715
n = 2,748
AA
TT
AA
TT

### Figure S6

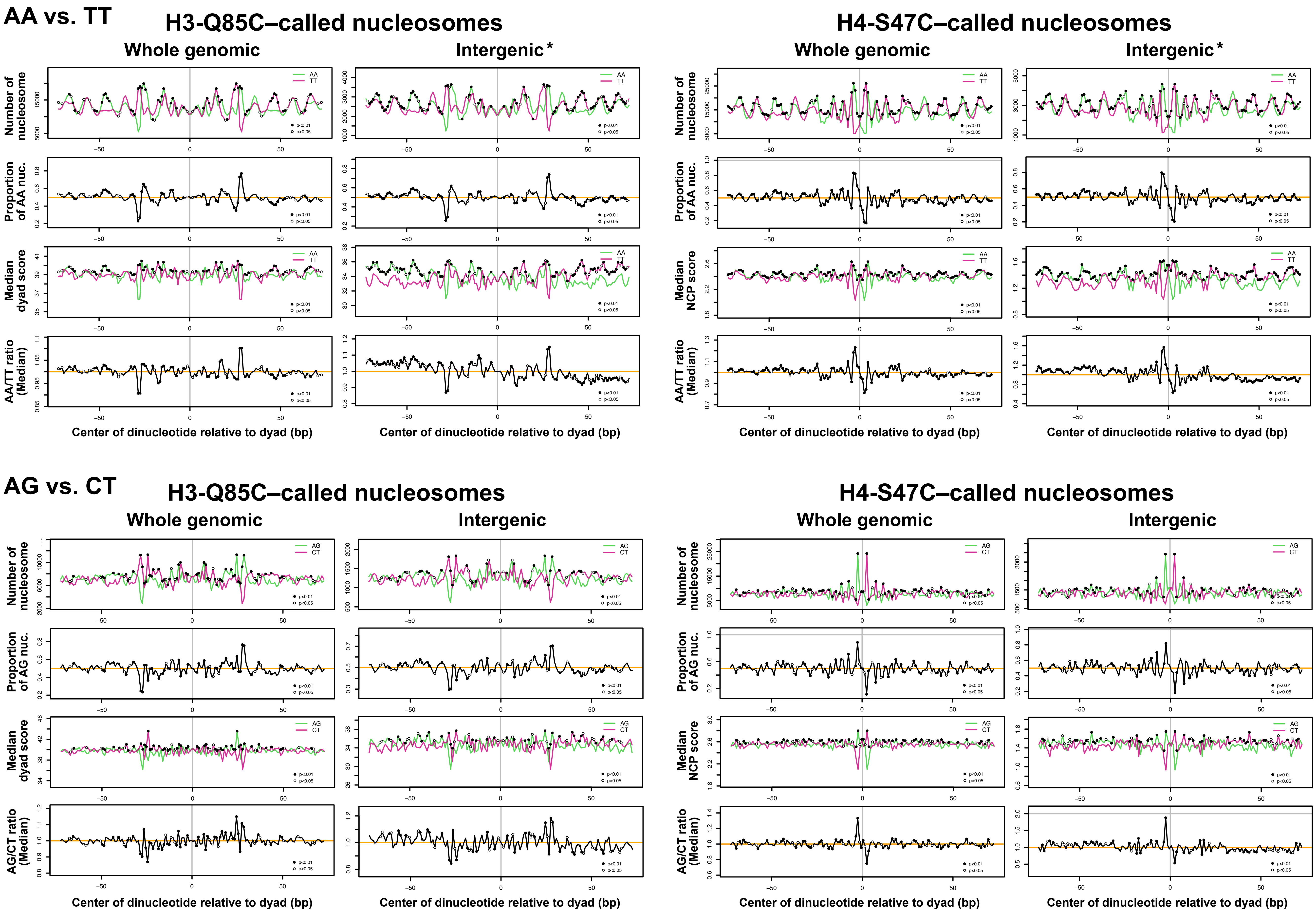
AA vs. TT
H3-Q85C–called nucleosomes
H4-S47C–called nucleosomes
Whole genomic
Intergenic*
Number of nucleosome
Proportion of AA nuc.
Median dyad score
Median NCP score
AA/TT ratio (Median)
AA
TT
p<0.01
p<0.05
Center of dinucleotide relative to dyad (bp)
AG vs. CT
H3-Q85C–called nucleosomes
H4-S47C–called nucleosomes
Whole genomic
Intergenic
Number of nucleosome
Proportion of AG nuc.
Median dyad score
Median NCP score
AG/CT ratio (Median)
AG
CT
p<0.01
p<0.05
Center of dinucleotide relative to dyad (bp)

### Figure S6, continued

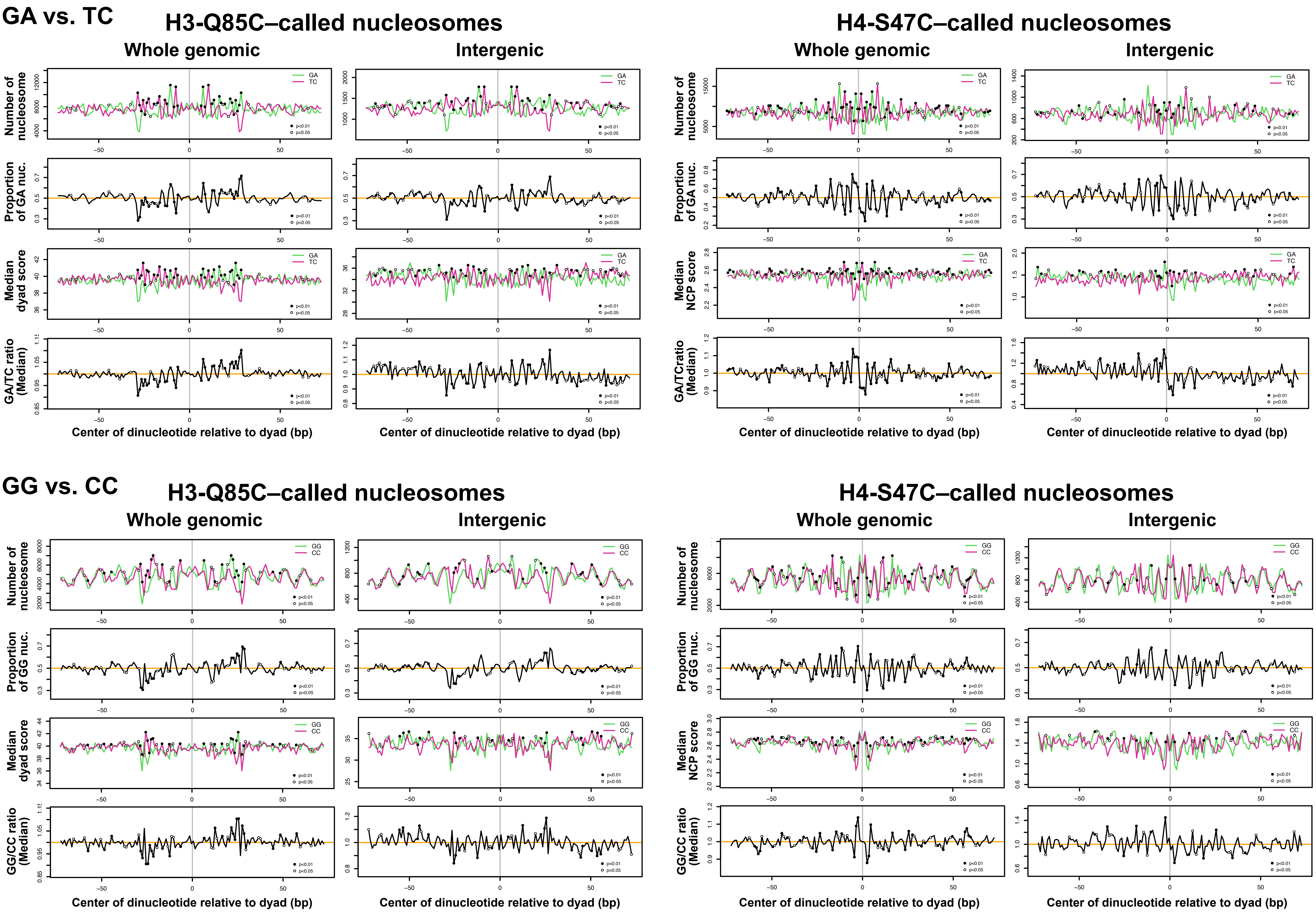

### Figure S7

## RR vs. YY

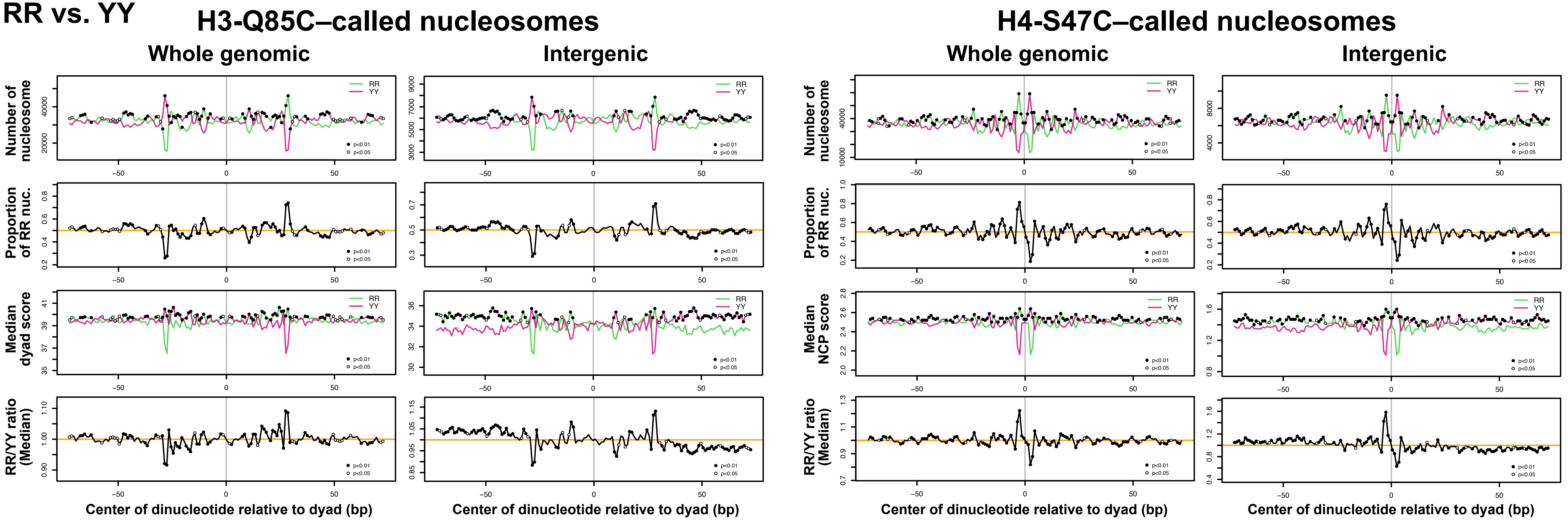

Figure S8

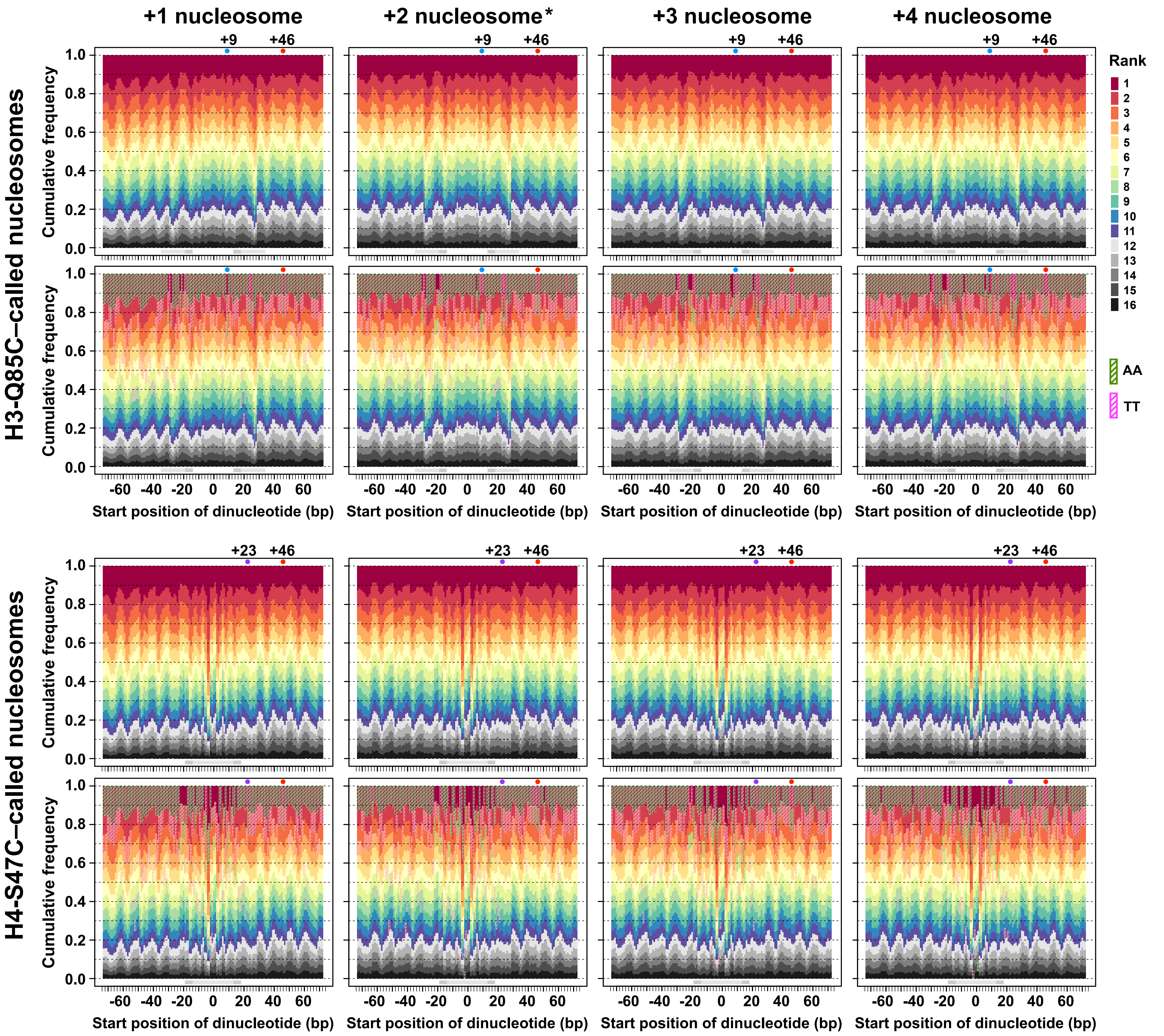

+1 nucleosome
+2 nucleosome*
+3 nucleosome
+4 nucleosome
+9
+46
+23
H3-Q85C–called nucleosomes
H4-S47C–called nucleosomes
Cumulative frequency
1.0
0.8
0.6
0.4
0.2
0.0
-60 -40 -20 0 20 40 60
Start position of dinucleotide (bp)
Rank
AA
TT
